# The RNase J-based RNA degradosome is compartmentalized in the gastric pathogen *Helicobacter pylori*

**DOI:** 10.1101/2020.05.11.085670

**Authors:** Alejandro Tejada-Arranz, Eloïse Galtier, Lamya El Mortaji, Evelyne Turlin, Dmitry Ershov, Hilde De Reuse

## Abstract

Post-transcriptional regulation is a major level of gene expression control in any cell. In bacteria, multiprotein machines called RNA degradosomes are central for RNA processing and degradation and some were reported to be compartmentalized inside these organelle-less cells. The minimal RNA degradosome of the important gastric pathogen *Helicobacter pylori* is composed of the essential ribonuclease RNase J and RhpA, its sole DEAD-box RNA helicase, and plays a major role in the regulation of mRNA decay and adaptation to gastric colonization. Here, the subcellular localization of the *H. pylori* RNA degradosome was investigated using cellular fractionation and both confocal and super-resolution microscopy. We established that RNase J and RhpA are peripheral inner membrane proteins and that this association was mediated neither by ribosomes, by RNA nor by the RNase Y membrane protein. In live *H. pylori* cells, we observed that fluorescent RNase J and RhpA protein fusions assemble into non-polar foci. We identified factors that regulate the formation of these foci without affecting the degradosome membrane association. Flotillin, a bacterial membrane scaffolding protein, and free RNA promote foci formation in *H. pylori*. Finally, RNase J-GFP molecules and foci in cells were quantified by 3D-single-molecule fluorescence localization microscopy. The number and size of the RNase J foci were found to be scaled with growth phase and cell volume as was previously reported for eukaryotic ribonucleoprotein granules. In conclusion, we propose that membrane compartmentalization and the regulated clustering of RNase J-based degradosome hubs represent important levels of control of their activity and specificity.

**Importance:** *Helicobacter pylori* is a bacterial pathogen that chronically colonizes the stomach of half of the human population worldwide. Infection by *H. pylori* can lead to the development of gastric pathologies such as ulcers and adenocarcinoma, that causes up to 800.000 deaths in the world each year. Persistent colonization by *H. pylori* relies on regulation of the expression of adaptation-related genes. One major level of such control is post-transcriptional regulation that, in *H. pylori*, largely relies on a multi-protein molecular machine, an RNA-degradosome, that we previously discovered. In this study, we established that the two protein partners of this machine are associated to the membrane of *H. pylori*. Using cutting-edge microscopy, we showed that these complexes assemble into hubs whose formation is regulated by free RNA and scaled with bacterial size and growth phase. Cellular compartmentalization of molecular machines into hubs emerges as an important regulatory level in the organelle-less bacteria.

## Introduction

Post-transcriptional regulation is one of the most important levels of control of gene expression in every kingdom of life. Ribonucleases (RNases) are key enzymes in post-transcriptional regulation, involved in RNA maturation and degradation. RNases often act in multi-protein complexes that are designated exosomes in Eukarya and Archaea and RNA degradosomes in bacteria and chloroplasts [for a review, see (1)]. RNA-degradosomes were established in several bacterial species and are defined by two core components, an RNase and an RNA helicase (2). These RNA helicases belong to the DEAD-box family and act by unwinding RNA, thereby allowing access of the ribonucleases to some of their target sites on RNAs. RNA degradosomes are widespread and vary in composition, although only few have been described in detail (1). Most RNA degradosomes reported so far are assembled on the essential endoribonuclease RNase E, like in *Escherichia coli, Caulobacter crescentus* or *Mycobacterium tuberculosis* (3–5). In the *E. coli* degradosome, RNase E serves as a scaffold for the binding of the DEAD-box RNA helicase RhlB, the metabolic enzyme enolase and the 3’-5’ exoribonuclease PNPase (2, 6). Nevertheless, our recent analysis on a representative set of 1,535 bacterial genomes revealed that RNase E is absent from about half of the bacterial species (1). Most of the remaining bacteria (47%), that lack RNase E, have either RNase Y or RNase J enzymes or both. RNase J and RNase Y, first identified in *Bacillus subtilis*, both display endoribonucleolytic activities but RNase J acts in addition as a 5’-3’ exoribonuclease (7, 8). Although being unrelated proteins, these enzymes constitute functional homologues of RNase E. In the Gram-positive bacteria *Staphylococcus aureus* and *B. subtilis*, RNA degradosomes comprising RNase Y and RNase J have been proposed, but to date their functionality was not clearly established and in *B. subtilis*, the complex was only detected after cross-linking (9).

In the Gram-negative pathogen *Helicobacter pylori*, we recently demonstrated the existence of a minimal RNA degradosome composed of two partners (10–12). *H. pylori* is a spiral shaped bacterium that persistently colonizes the stomach of half of the human population worldwide. Infection by this pathogen causes chronic gastritis, peptic ulcers, MALT lymphoma and gastric carcinoma, which is responsible for 800,000 deaths per year in the world (13, 14). Persistent colonization of the hostile acidic gastric niche by *H. pylori* is associated with the development of severe pathologies such as gastric cancer. *H. pylori* possesses a small genome (1.6 Mb) and a reduced number of transcriptional regulators. Therefore, post-transcriptional regulation has been proposed to play a major role in the adaptive response of *H. pylori* (15).

The RNA degradosome of *H. pylori* is composed of the essential RNase J protein and of RhpA, the sole DEAD-box RNA helicase of this bacterium (10–12). *In vitro*, each protein of this complex stimulates the activity of the other, thus demonstrating that they form a functional RNA degradosome (10). Moreover, using sucrose gradients, both proteins were detected in association with translating ribosomes, suggesting a coupling between translation and RNA degradation (10). More recently, we performed an RNAseq analysis of an *H. pylori* RNase J-depleted strain and observed that the amounts of 55% of mRNAs and 40% of antisense RNAs (asRNAs) are increased more than 4-fold in the mutant in comparison with the parental wild type strain (11). Only 12% of intergenic non-coding RNAs are affected by RNase J depletion, suggesting that they are poor targets for this enzyme. Taken together, these results indicate a major role of RNase J in mRNA decay in *H. pylori*.

The ribonuclease activities of the RNA degradosomes are generally essential for bacterial survival but they are also potentially destructive if these molecular machines degrade RNAs that are important for growth. Accordingly, several levels of control of degradosomes have been identified (1). One of them is compartmentalization, whose importance in the organelle-less prokaryotes is increasingly being recognized. Compartmentalization of metabolic processes is essential for any cell in order to optimize enzymatic reactions and avoid unwanted effects. Emerging structures that can be compartmentalized are associated to membranes or are the so-called membrane-less organelles formed by liquid-liquid phase separation (LLPS). Some bacterial RNA degradosomes have recently been reported to be compartmentalized (1). The RNase E-based degradosome of *E. coli* localizes at the inner membrane of the cell (16, 17) through a short amphipathic helix present at the C-terminal half of RNase E (*Eco*RNase E). Deletion of the *Eco*RNase E membrane anchor does not result in massive transcriptomic changes but in a global slowdown of RNA degradation (18), although another report suggests that the *Eco*RNase E membrane anchor is responsible for the shorter average half-life of membrane protein-encoding transcripts (19). RhlB, the helicase partner of *Eco*RNase E displays the same localization and its membrane targeting depends on RNase E (20). In addition, *Eco*RNase E fluorescent fusion proteins form short-lived foci and rapidly diffuse across the inner membrane (16, 20). Interestingly, similarly to the situation in *H. pylori*, the *E. coli* degradosome can form stable complexes with translating ribosomes (21). In *C. crescentus*, RNase E (*Cc*RNase E) lacks a membrane anchor and has been shown to be cytoplasmic (22). *Cc*RNase E-YFP fusion proteins form clusters in the cell that are dynamically assembled, change with cell cycle or stress exposure and are proposed to be the sites where RNA cleavage occurs (23, 24). These foci present characteristics of LLPS similar to eukaryotic messenger ribonucleoprotein (mRNP) granules, such as p-bodies or stress granules that segregate different RNA molecules from the cytoplasm and can control their fate (1).

In *S. aureus* and *B. subtilis*, the proposed RNA degradosome relies on RNase Y, which is inserted at the inner membrane through its N-terminal transmembrane region (25–27). In *S. aureus*, FloA, a membrane scaffolding protein homologous to eukaryotic flotillin promotes the oligomerization and the activity of RNase Y (28). In *B. subtilis*, a homogeneous membrane localization of an RNase Y-GFP fusion was observed (26), and recently found to be dynamic and form foci similar to those of RNase E (29). Under conditions where RNase Y was less active, the foci were more abundant and increase in size suggesting that these structures represent a less active form of the enzyme (29). RNases J1 and J2 were found to localize in the cytoplasm and were excluded from the nucleoid (26). Analysis of GFP-fusions of the other proposed partners of the *B. subtilis* RNA-degradosome (PNPase, enolase, phosphofructokinase and the DEAD-box RNA helicase CshA) suggested that they are mainly cytoplasmic (26).

Thus, compartmentalization seems to be a frequent feature of bacterial RNA degradomes that is submitted to regulation and that probably controls its activity. Therefore, the subcellular localization of RNA degradosomes is an important question that was only addressed for a few degradosomes. Here, this question was addressed using the major pathogen *H. pylori* as a model organism in which the minimal RNase J-based degradosome plays a central role in post-transcriptional regulation.

In this report, we show that both *H. pylori* RNase J and RhpA target the inner membrane and form foci. We demonstrated that RNase J foci are subject to variations as a function of growth phase and antibiotic exposure, suggesting differential regulation of the activity of the RNA degradosome between foci-forming and non-foci-forming forms.

## Results

### Localization of RNase J and RhpA at the *H. pylori* inner membrane

The subcellular localization of RNase J and RhpA was determined in *H. pylori* strain B128 using a cellular fractionation protocol adapted to this bacterium [after (30)]. Separation of the different fractions was validated using control antibodies, anti-BabA for the outer membrane (OM), anti-MotB for the inner membrane (IM) and anti-AmiE for the soluble extract (SE) (Fig. S1A). Western blots using specific anti-RNase J and anti-RhpA antibodies (10) revealed that both proteins are mostly present at the inner membrane, with a small amount in the soluble fraction (Fig. 1A). No protein was detected in the outer membrane. The same result was obtained with another *H. pylori* strain, the first sequenced well studied 26695 strain (Fig. S1B).

**Figure 1.**
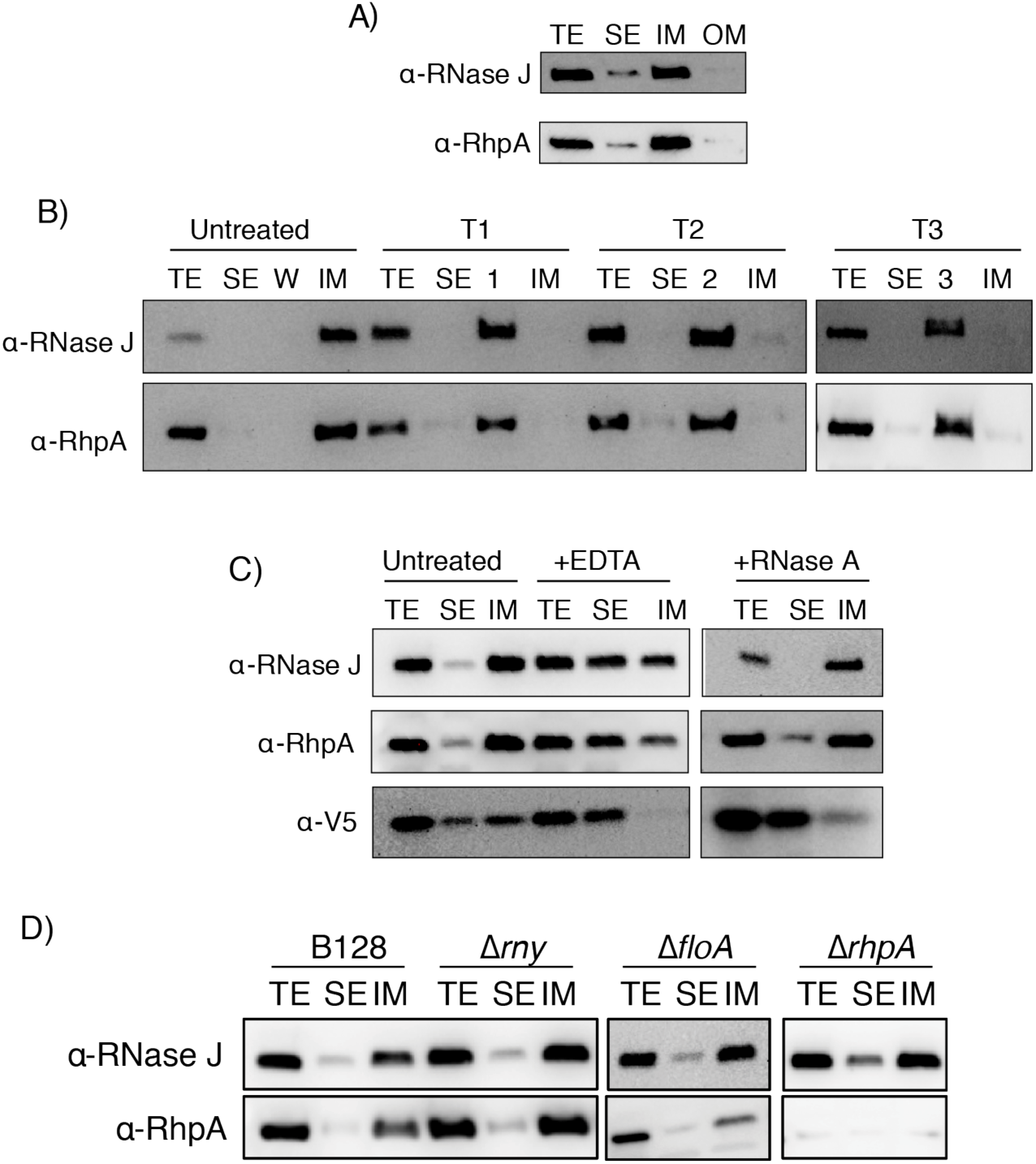
The two partners of the RNA degradosome of *H. pylori*, RNase J and RhpA, are associated with the inner membrane in a peripheral manner, independently of ribosomes and RNA and this association does not depend on RNase Y, flotillin nor RhpA. Cellular compartments of *H. pylori* B128 strain were separated by fractionation; each sample corresponds to the same initial number of bacteria. Experiments were performed in triplicate. TE: total extract fraction, SE: soluble extract fraction, IM: inner membrane fraction, OM: outer membrane fraction. A) Western blot with antibodies against RNase J and RhpA on the different cellular fractions of WT *H. pylori* strain B128. B) Western blot with antibodies against RNase J and RhpA on the different cellular fractions of WT *H. pylori* upon treatment of the membrane fraction with 6M urea (T1), 100 mM Na_2_CO_3_ (T2) or 2M NaCl (T3), untreated fractions are shown as a control. Labels 1, 2 and 3 indicate the wash fractions with their respective treatments T1, T2 or T3. C) Western blot with antibodies against RNase J, RhpA and a V5 tag (that marks the L9 ribosomal protein) on the different cellular fractions of an L9-V5-expressing *H. pylori* strain, under untreated conditions and upon treatment with 20 mM EDTA or 1 µg/mL RNase A. D) Western blot with antibodies against RNase J and RhpA on the different subcellular fractions of WT, Δ*rny*, Δ*floA* or Δ*rhpA H. pylori* strains.

### RNase J and RhpA are peripheral membrane proteins

Since no transmembrane domains are predicted in the sequences of RNase J and RhpA (Fig. S1CD), we investigated the nature of the association of the RNA degradosome to the membrane. For this purpose, different treatments were applied to the purified membrane fractions, which were subsequently subjected to ultracentrifugation to separate the supernatant, containing the extracted peripheral membrane proteins, from the pellet that contains the integral membrane proteins. These treatments include (i) 6 M urea, a chaotropic agent that weakens the hydrophobic interactions without disrupting the lipid bilayer, (ii) alkaline pH (100 mM Na_2_CO_3_ pH 11) and (iii) high salt concentration (2M NaCl). The three treatments consistently dissociated both RhpA and RNase J from the pelleted membrane fraction (Fig. 1B), clearly indicating that these two proteins are associated to the membrane through electrostatic and hydrophobic interactions and are thus peripheral inner membrane proteins in *H. pylori*.

### Membrane targeting of the degradosome is independent of ribosomes and RNA

Our previous work revealed that at least a fraction of the *H. pylori* degradosome proteins was associated with translating ribosomes (10). To examine whether ribosomes might have been co-purified with the inner membrane fraction, and/or whether the RNase J and RhpA membrane association could be mediated by ribosomes, we constructed an *H. pylori* strain expressing a fusion between the L9 ribosomal protein of the large subunit and a V5 epitope from the native locus at the chromosome (Fig. S2A), that we used as a reporter for ribosome localization. Using this strain, we observed that, upon fractionation, the ribosomes were approximately evenly distributed between the inner membrane and soluble fractions (Fig. 1C). Treatment with EDTA, that through Mg^2+^ chelation causes dissociation of the ribosomal subunits, results in a complete delocalization of L9-V5 from the membrane fraction. EDTA treatment only partially delocalized RNase J and RhpA to the soluble extract (Fig. 1C). Given that EDTA not only acts on ribosomes but also on the divalent cation pool of the cell and consequently on electrostatic interactions, it is difficult to dissociate the two effects in the analysis of this localization. To clarify this effect, we performed a treatment with RNase A, which degrades all unprotected RNAs, and therefore dissociates polysomes. Indeed, this treatment detaches most of the ribosomes from the inner membrane. Importantly, under these conditions, neither RNase J nor RhpA are delocalized towards the soluble extract, which strongly suggests that their membrane interaction is not mediated by ribosomes nor RNA and that it is, at least in part, cation-dependent (Fig. 1C). Thus, ribosomes and RNA are not required for the inner membrane localization of the two proteins of the *H. pylori* degradosome and divalent cations are most probably important for their electrostatic membrane binding.

### Membrane targeting of the degradosome is independent of RNase Y and flotillin

We wanted to explore the possibility that RNase J and RhpA could be indirectly bound to the membrane through another partner. In *B. subtilis*, RNase J1 was reported to be associated with the membrane-bound RNase Y. However, *B. subtilis* RNase J1 main localization seems to be cytoplasmic (26). A transmembrane domain is also predicted in the sequence of the *H. pylori* RNase Y protein (Fig. S1E). Therefore, an *H. pylori* mutant carrying a deletion of the corresponding *rny* gene was constructed and used to examine the impact of its absence on the localization of the degradosome. As shown in Fig. 1D, in this mutant both RNase J and RhpA are still associated with the inner membrane, indicating that RNase Y is not the membrane anchor of the degradosome in *H. pylori*.

It has also been shown that flotillin, a bacterial membrane-scaffolding protein related to the flotillins from eukaryotic lipid rafts, plays a role in the oligomerization state of the *S. aureus* RNase Y, a component of the RNA degradosome in this organism (28). *H. pylori* possesses only one flotillin that was recently identified (31). The corresponding *floA* gene (HPB128_21g23 in strain B128) is located immediately downstream of the *rhpA* gene (Fig. S2A). In order to test whether the membrane localization of RNase J and/or RhpA was dependent on the presence of flotillin, we constructed a Δ*floA H. pylori* mutant and examined the membrane localization of RNase J and RhpA. We found that the inner membrane association of these proteins was not affected in the Δ*floA* mutant as compared to the wild type strain and thus that flotillin is not required for their targeting to the membrane (Fig. 1D).

We also wanted to test the respective role of each protein in the membrane localization of the other partner. Since RNase J is an essential protein, only a *ΔrhpA* mutant could be examined. We observed that RhpA is not required for the membrane targeting of RNase J. Under these conditions, the larger amount of RNase J in the soluble extract was attributed to its overexpression in the *ΔrhpA* mutant that we previously reported (12) (Fig. 1D).

### RNase J-GFP and RhpA-CFP form foci in live *H. pylori* cells

We then wanted to analyze the localization of RNase J and RhpA by confocal fluorescence microscopy in live *H. pylori* cells. For this, we constructed *H. pylori* B128 strains that express RNase J fused in C-terminus to the GFPmut2 fluorescent protein or RhpA fused in C-terminus to the SCFP fluorescent protein. The *rnj-gfp* and *rhpA-cfp* fusion genes were introduced in the chromosome and expressed from their native promoters, replacing the original copies of *rnj* or *rhpA*, respectively (Fig. S2A). Adding a tag to the C-terminus of RNase J and RhpA does not seem to prevent their functions, as both fusion strains show no growth defect, taking into account that RNase J is an essential gene, and that the Δ*rhpA* mutant has a considerable growth defect (11, 12). The strains carrying these fusions preserve the characteristic spiral shape and no morphological defects as visualized by phase contrast microscopy (Fig. S2B). Western blots with anti-RNase J and anti-RhpA antibodies showed that the fusion proteins were well expressed and did not undergo *in vivo* degradation (Fig. S2C).

By cellular fractionation, we found that the RNase J-GFP fusion membrane localization was similar to that of native RNase J (Fig. S2C). In contrast, we observed that the RhpA C-terminal CFP fusion causes a partial delocalization of RhpA towards the soluble extract when compared to the native RhpA protein (a similar delocalization was observed with a RhpA N-terminal CFP fusion).

Confocal fluorescence microscopy revealed that RNase J-GFP is visible as intense discrete foci that lie at the periphery of live bacteria (Fig. 2A). In the case of RhpA-CFP, spots were also detected at the periphery of the bacteria, although with more background throughout the cell (Fig. 2A). This is probably caused by the partial membrane delocalization of this fusion as mentioned above.

**Figure 2.**
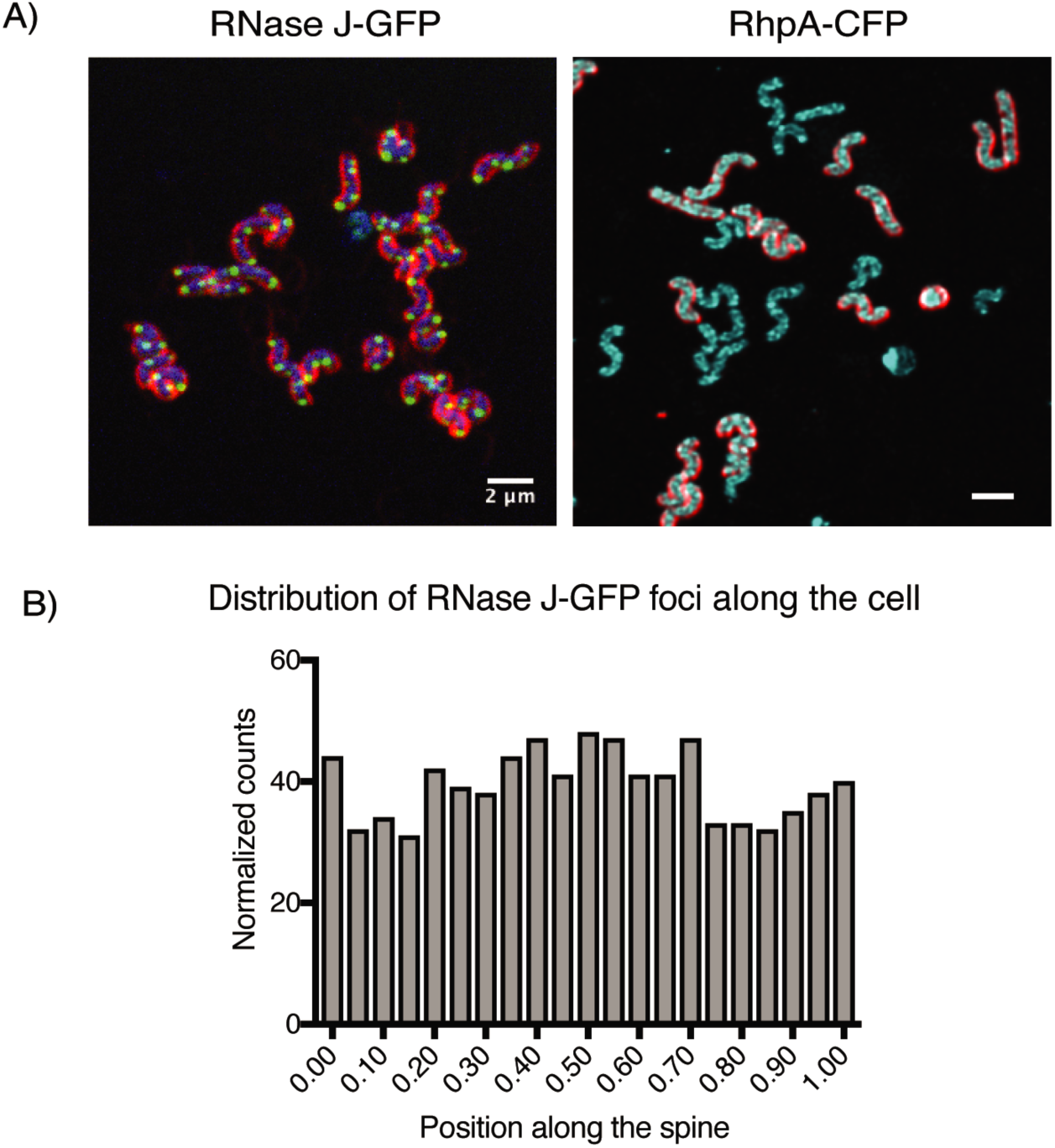
RNase J and RhpA form foci inside the cells that do not have a polar localization. A) Representative composite confocal microscopy images of live *H. pylori* cells expressing RNase J-GFP (green) or RhpA-CFP (cyan). In the RNase J-GFP image, blue is DNA (Hoescht 33342) and, in both images, red are membranes (FM4-64). Experiments were performed at least 3 times. B) Histogram showing the number of foci of RNase J-GFP that are located in each position along the spine of the *H. pylori* cells.

There are very few reports on GFP fusions in live *H. pylori* and none with CFP. Therefore, several controls were performed to ensure that the degradosome foci were not generated by self-aggregation of GFPmut2 and SCFP fusions in *H. pylori. H. pylori* over-expressing from a plasmid either GFPmut2 and SCFP alone displayed no foci but rather an intense diffuse fluorescence signal distributed all over the cells (Fig. S2D). In addition, immunofluorescence using anti-FLAG antibodies on fixed *H. pylori* cells expressing RNase J-FLAG or RhpA-FLAG fusions from the chromosome revealed foci similar to those of live cells (Fig. S2E).

We then analyzed whether the RNase J-GFP foci localize to a preferential site in the cell, such as the poles. As a first step to measure the amount of polar foci, the nucleoid signal was used to assign each detected focus to the corresponding parental cell. Then, a central axis for the cells was established and the foci were assigned to the point nearest to them within the axis, in such a way that if a focus was assigned to the extremes of the axis, it would be classified as a polar focus while if assigned to the middle of the axis, it would be classified as septal. With a total of 1,403 foci analyzed, we found that they are randomly distributed in the cells and that only 5% of the total foci were located at the poles (Fig. 2B).

The foci formed by the RNase E-degradosomes, either membrane associated as in *E. coli* or cytoplasmic as in *C. crescentus* were reported to be dynamically assembled (1). Thus, the position of the RNase J-associated degradosome foci was monitored in live *H. pylori* cells every 6 seconds during 3 min and compared with those of fixed cells. No significant movement of the foci was detected suggesting that the RNase J-based foci are static in *H. pylori* (Fig. S2F and supplementary videos 1-2).

Our data show that the proteins of the *H. pylori* degradosome form foci that are likely physiologically compartmentalized structures. RNase-J foci analysis revealed that they are (i) static and (ii) non-polar but rather randomly distributed along the cell.

### Growth phase and antibiotic treatment impact the formation of RNase J-GFP foci

As we showed that the RNase J-GFP fusion protein localizes like the native RNase J, we decided to use this fusion to further investigate what factors might influence foci formation. First, we quantified the number of foci per cell by automatically detecting the foci particles and assigning them to the nearest nucleoid. Every measure was normalized to the area of each nucleoid in order to avoid an overestimation of the number of foci due to dividing cells. The median number of foci per nucleoid was about 3 during exponential growth phase with a range between 0 and 6 (Fig. 3AB).

**Figure 3.**
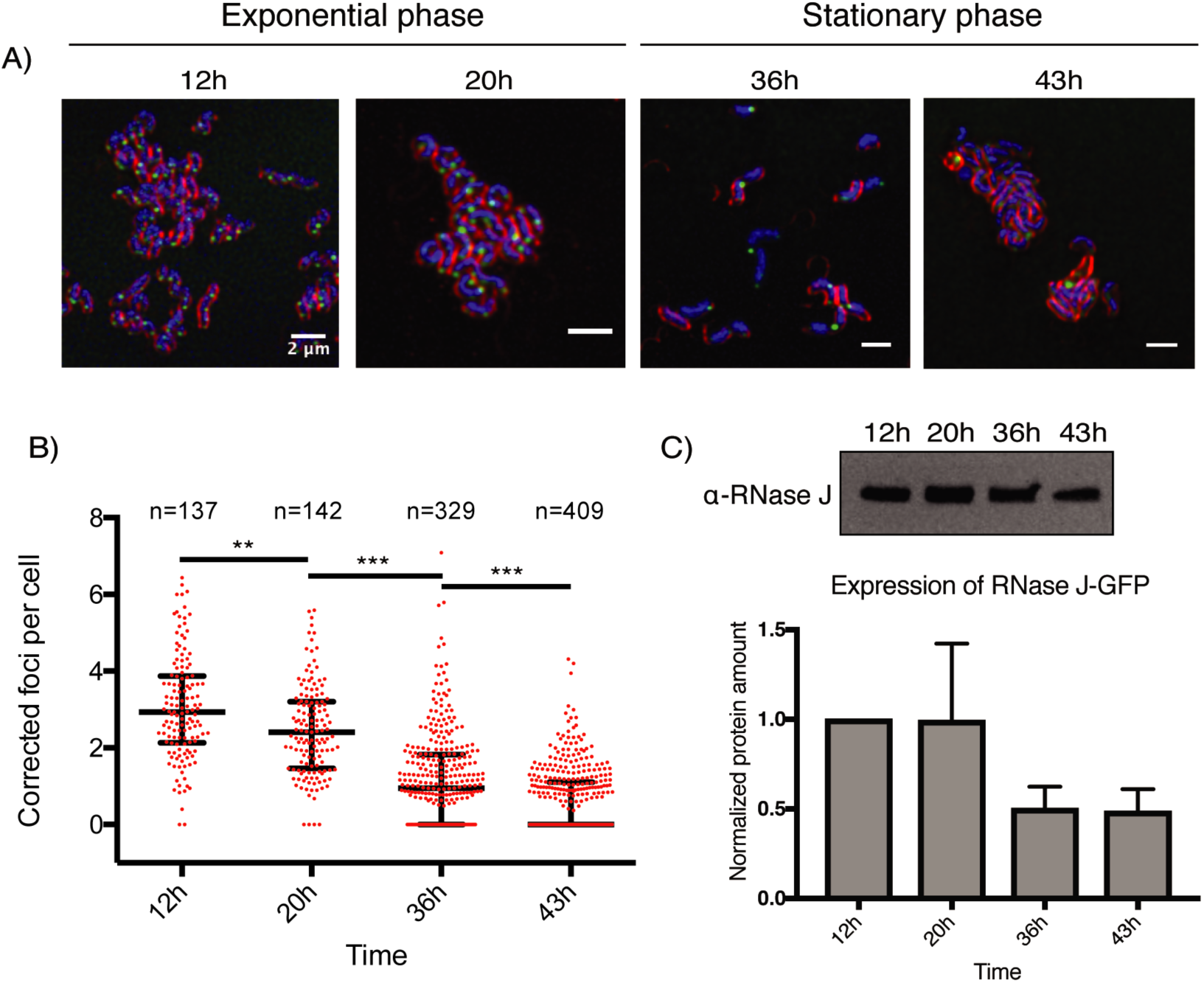
The number of RNase J-GFP foci per cell is progressively reduced along the *H. pylori* growth curve. A) Representative composite confocal microscopy images of the RNase J-GFP expressing strain at different time points (in hours) along the growth curve. In blue, DNA (labeled with Hoechst 33342); in green, RNase J-GFP; and in red, the membrane (labeled with FM4-64). The experiment was performed in triplicate. B) Quantification of the amount of foci per cell, normalized by the nucleoid area, along the growth curve. “n” corresponds to the number of cells analyzed for each condition. The median value is represented by a horizontal bar, and the error bars correspond to the interquartile range. *p<0.05, ** p<0.005, ***p<0.0005. C) Western blot (upper panel) and quantification (lower panel) of RNase J-GFP of cells taken at different time points along the growth curve and normalized on total proteins. The experiments were reproduced twice. The differences are not statistically significant, p-value of 0.11.

Next, we wanted to check whether there was an effect of the growth phase on the amount of foci per cell. At the time we took *H. pylori* bacteria in stationary phase, all cells had a spiral shape. We observed that the median number of detected RNase J-GFP foci is steadily reduced throughout the growth phase, with a median of 2.4 foci/cell at 20h (exponential phase cells), 0.95 foci/cell at 36h and 0 foci/cell at 43h (stationary phase cells) (Fig. 3AB). Cell fractionation and Western blots using anti-RNase J antibodies indicated that the levels of RNase J-GFP are indeed slightly reduced in stationary phase, but not sufficiently to explain the disappearance of the foci (Fig. 3C). In eukaryotes, p-bodies and stress granules require untranslated mRNAs to assemble. Similarly, formation of the RNase E cytoplasmic foci (BR bodies) in *C. crescentus* relies on free mRNAs (23). Therefore, we tested whether 30 min treatments with antibiotics known to affect the amount of translating or ribosome-free mRNAs in cells alter the number of foci in exponentially growing *H. pylori* bacteria. No effect on nucleoid condensation was observed under these conditions. The antibiotics were rifampicin that inhibits transcription and, as a consequence, causes a decrease in the total amount of mRNAs and chloramphenicol that blocks ribosomes on the mRNAs that they are translating, reducing the pool of ribosome-free mRNAs. Upon treatment with rifampicin and chloramphenicol, we found a significant decrease in the median amount of foci per cell that was more pronounced in the case of rifampicin (63% reduction) than with chloramphenicol (36% reduction) (Fig. 4A) when compared with untreated cells. Finally, no significant difference in the number of foci was observed after puromycin treatment, an antibiotic that leads to the dissociation of the ribosomal subunits, increasing the amounts of untranslated cytoplasmic mRNAs.

**Figure 4.**
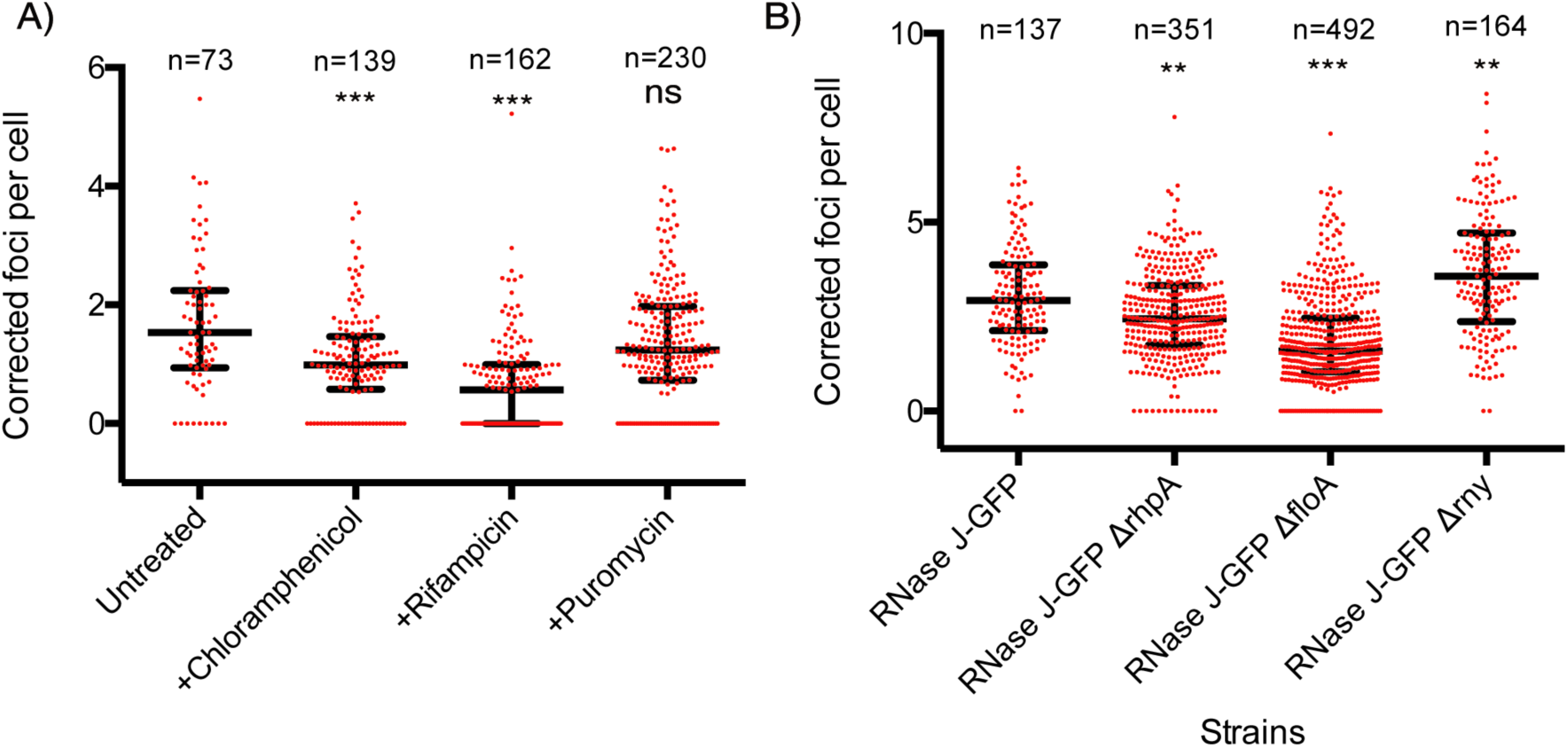
The number of RNase J-GFP foci per cell is affected by antibiotics and in different mutants. “n” corresponds to the number of cells analyzed for each condition. The median value is represented by a horizontal bar, and the error bars correspond to the interquartile range. Experiments were performed in triplicate. *p<0.05, **p<0.005, ***p<0.0005, “ns” is non-significant. A) Quantification of the number of RNase J-GFP foci, normalized by the nucleoid area, in untreated cells and upon treatment with chloramphenicol, rifampicin or puromycin. B) Quantification of the number of RNase J-GFP foci, normalized by the nucleoid area, in wild-type cells and in bacteria deleted for the genes encoding RNase Y, flotillin or RhpA.

Furthermore, we examined whether the membrane localization of RNase J was perturbed by either the growth phase or the antibiotic treatments and we found that it remains associated to the inner membrane (Fig. S3AB).

These data suggest that ribosome-free mRNAs are among the factors promoting foci formation in *H. pylori* and that the formation of foci is not only correlated with the amount of RNase J-GFP.

### Flotillin and RNase Y influence the number of RNase J-GFP foci

As shown above, RhpA, RNase Y and flotillin did not significantly alter the membrane localization of RNase J in *H. pylori*. However, we wondered whether these proteins might impact the formation of foci. Therefore, Δ*rny*, Δ*flot* and Δ*rhpA* mutations were introduced in the strain expressing RNase J-GFP and the number of foci per cell was quantified in exponential phase. We verified that the membrane localization of RNase J-GFP fusion was not modified in these mutants (Fig. S3C) and that the non-polar distribution of the foci was conserved.

We found that the median number of foci was significantly increased (by 22%) in the Δ*rny* strain (Fig. 4B). In both the Δ*rhpA* and Δ*floA* mutants, the amount of foci was found to be reduced (by 17 and 46%, respectively), with the effect being much stronger in the flotillin-deficient mutant (Fig. 4B). Taken together, these results indicate that the number of foci per cell is not linearly correlated to the amounts of RNase J-GFP protein and that additional factors determine their formation such as the growth phase, flotillin and RNase Y.

### Super-resolution microscopy to quantify the RNase J-based degradosome foci

We decided to explore in more details the properties of the RNase J degradosome foci in *H. pylori* both in exponential and stationary phase. Therefore, we applied, for the first time in this bacterium, direct STochastic Optical Reconstruction Microscopy, dSTORM, a powerful tool that allows a 15 nm location precision of individual molecules in X, Y and Z axes. For this analysis, *H. pylori* was fixed and permeabilized, membranes were labeled with the wheat germ agglutinin (WGA) lectin coupled to Alexa Fluor 555, and GFP was labeled with anti-GFP nanobodies coupled to sulfocyanin 5 (Cy5), allowing us to analyze the localization of RNase J-GFP at the single molecule level.

First, we observed that the foci of RNase J-GFP are indeed located at the membrane in both exponential and stationary phase (Fig. 5 and supplementary videos 3-6). In addition, higher sensitivity of super-resolution microscopy allowed us to visualize more foci than conventional confocal microscopy. A median number of 4 foci per cell was observed in exponential phase and 2 in stationary phase (Fig. 6A), in agreement with the reduction of foci seen by confocal microscopy. Furthermore, we could quantify the number of RNase J-GFP molecules per focus and analyze their dimensions in both phases. In exponential phase, we measured a median of 307 RNase J-GFP molecules per cell (Fig. 6B), with a median of 19 RNase J-GFP molecules in each focus (Fig. 6C) that present a median volume of 5.91e-4 μm^3^ (Fig. 6D). In stationary phase, we detected a median of 104 total molecules per cell (Fig. 6B), the foci contain a median of 8 molecules (Fig. 6C) and present a volume of 6.75e-5 μm^3^ (Fig. 6D), approximately nine times less than the foci in exponential phase. Therefore, foci are smaller and contain less RNase J-GFP molecules in stationary phase. As expected, the cell volume also changes during growth, a median volume of 0.7 µm^3^ was measured in exponential phase that changed to 0.36 µm^3^ in stationary phase (Fig. 6E). If we correct by the measured bacterial volume, the concentration of RNase J molecules per cell is marginally changing between exponential and stationary phase and so is the number of foci per cell, whereas the foci volume and the amount of RNase J per foci suffer more important changes. Therefore, we can conclude that the amount and size of the RNase J foci are scaled with the cell volume and thus with growth stage.

**Figure 5.**
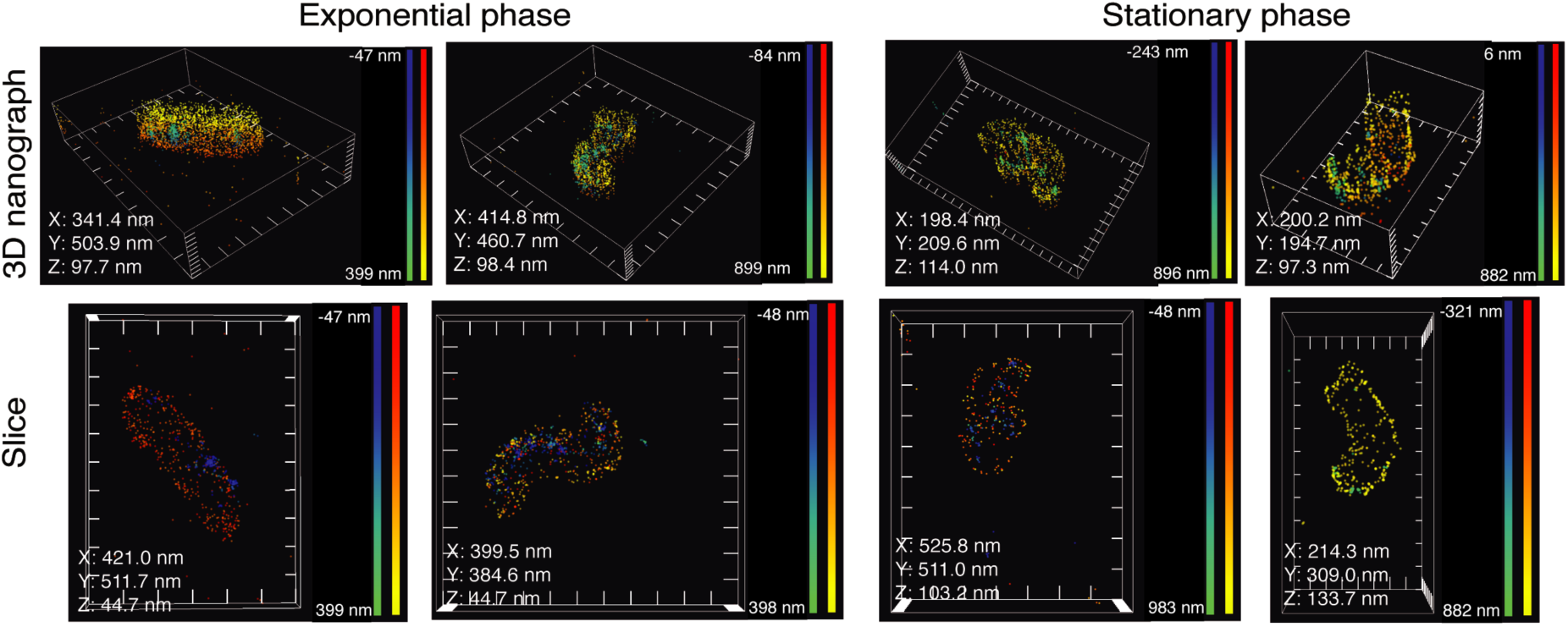
Visualization of the foci of RNase J-GFP in exponential and stationary phase by dSTORM super-resolution microscopy. Representative single-molecule localization microscopy images showing the membrane of *H. pylori* as visualized after labeling with WGA-AF555 (yellow-red) and the RNase J-GFP foci after labeling with anti-GFP-Cy5 nanobodies (blue-green). The upper panels show the full 3D volume of representative cells and the lower panels show a 2D longitudinal slice of the bacteria. The X, Y and Z values in each image indicate the distance between the ticks of the respective axes in the picture. The color gradients red-yellow (for the WGA-AF555) and blue-green (for anti-GFP-Cy5 nanobodies) indicate the distance of each dot with respect to the coverslip, as indicated in the corresponding scale bars in each panel. The experiment was performed 3 times.

**Figure 6.**
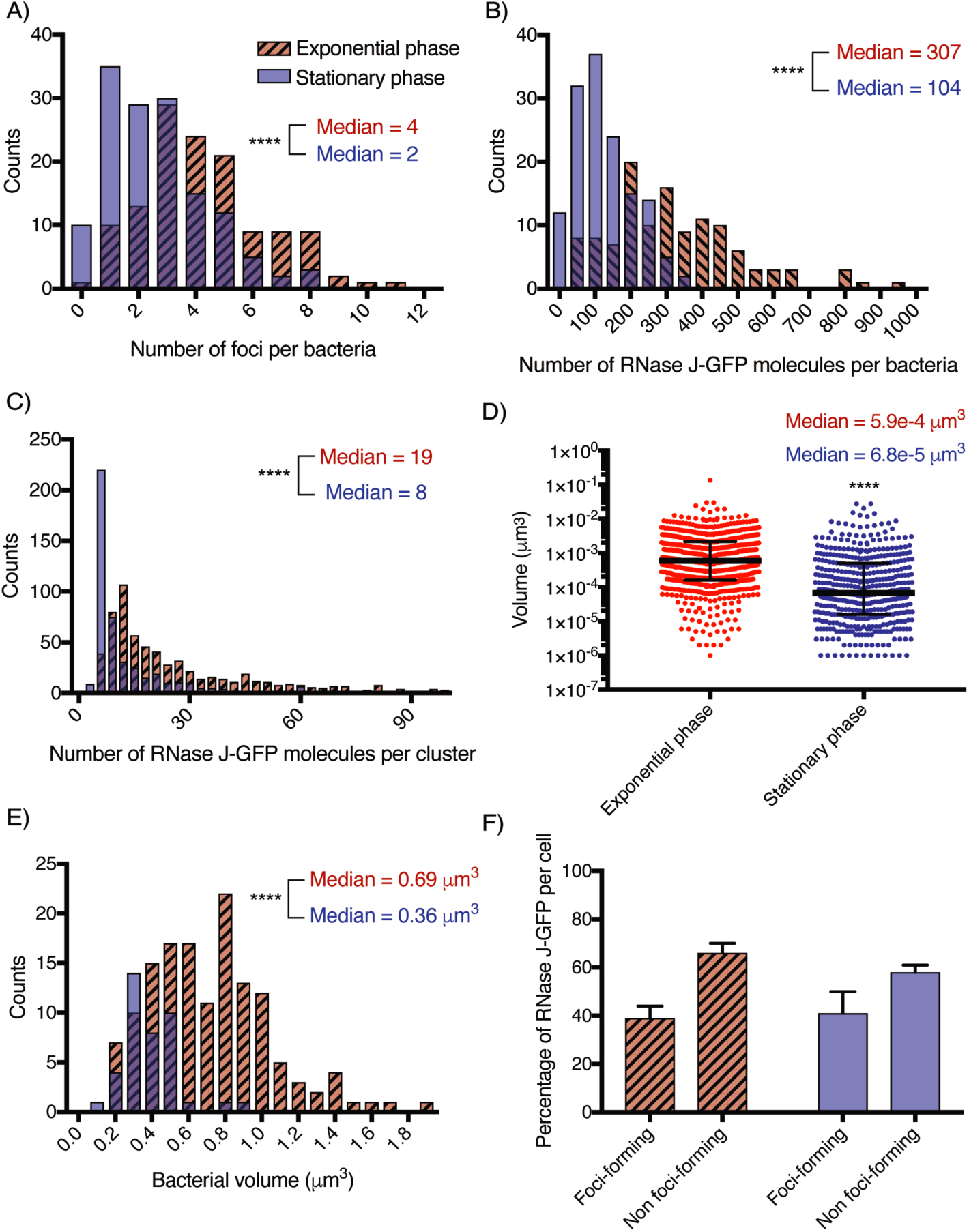
dSTORM quantification of the RNase J-GFP foci of *H. pylori* cells in exponential (red hatched bars) and stationary phase (blue bars). The experiment was carried out 3 times and correspond to that of Figure 5. A) Distribution histogram of the amount of foci per cell in exponential (n=121 cells) and stationary phase cells (n=191 cells). Median values under both conditions are indicated. B) Distribution histogram of the number of RNase J-GFP molecules per cell in exponential and stationary phase cells. Median values under both conditions are indicated. C) Distribution histogram of the number of RNase J-GFP molecules per cluster in exponential and stationary phase cells (n=535 clusters in exponential phase and n=371 clusters in stationary phase). Median values under both conditions are indicated. D) Distribution histogram of the volume of the RNase J-GFP clusters in exponential and stationary phase cells. Median values under both conditions are indicated. E) Distribution histogram of the volume of *H. pylori* cells in exponential phase (n=142 cells) and stationary phase (n=40 cells). For the A, B, C, D and E panels, ****p<0.0001. F) Bar graph of the proportion of foci-forming and non-foci-forming RNase J-GFP molecules calculated on the mean values in exponential and stationary phase cells.

Finally, we also found that under both growth phases, the mean proportion of total RNase J-GFP molecules that are clustered in foci is the same (40%), irrespective of the total amount of RNase J-GFP (Fig. 6F). Thus, the foci are not the consequence of RNase J reaching a threshold concentration that causes them to spontaneously aggregate into foci but are rather formed by a regulated mechanism as suggested above. Furthermore, the fact that a significant fraction of RNase J-GFP (60%) does not form foci (Fig. 6F), suggests the existence of two different populations of RNase J that may play different physiological roles.

## Discussion

In this work, we uncovered the inner membrane association of the *H. pylori* RNA degradosome. RNase J and RhpA, the two protein partners of this minimal degradosome, present no predictable membrane targeting sequences; they interact with the inner membrane through hydrophobic and electrostatic interactions and are thus peripheral membrane proteins of *H. pylori*. This membrane localization of the RNA degradosome is independent of RNA and relies neither on the membrane-bound RNase Y nor on the sole flotillin of *H. pylori*. The association of the degradosome partners to the *H. pylori* membrane might, however, be mediated by another protein, yet to be identified. During our previous studies, RNase J and RhpA were detected in association with purified ribosomes (70S particles and polysomes) (10). The present results show that the membrane localization of the degradosome is not mediated by or dependent on ribosomes. Thus, two different possibilities arise: (i) the ribosomal-associated degradosome fraction corresponds to the small amount of proteins detected in the soluble fraction, which might play a function different from that of the membrane bound population, or (ii) the small ribosome-bound fraction of the degradosome is also attached to the membrane and plays a role only with ribosomes that are localized in the vicinity of the membrane.

Only a few studies reported the subcellular localization of RNA degradosomes in other bacteria. The localization varied depending on the organism and did not correlate with the nature of the scaffolding RNase it contains. In *E. coli*, the degradosome is targeted to the inner membrane through an amphipathic helix of RNase E. Strikingly, this degradosome also presents both inner membrane and ribosomal localizations (20, 21). We believe that this “evolutionary convergence” indicates that the dual localization of the RNA degradosome might have a functional significance. In contrast, RNase E of *C. crescentus* does not contain this helix and is cytoplasmic (22). Concerning the potential RNase Y-based degradosome of the Gram-positive bacterium *B. subtilis*, the results were rather contradictory. RNase Y is, as expected, membrane associated, but RNase J1, which is one of its proposed interacting partners, does not follow the same distribution, being cytoplasmic (26, 32). In contrast, in a recent work a fraction of RNase J2 of *Streptococcus mutans* was found to localize at the membrane (33). In *B. subtilis*, RNase Y was recently found to assemble at the membrane into dynamic short-lived foci that increase in number and size upon rifampicin treatment or deletion of the Y-complex, a protein complex that modulates RNase Y activity (29). Their data suggest that RNase Y foci represent, in contrast to those of RNase E, a less active form of the enzyme (29).

The differences in the localization of bacterial degradosomes might also be related to the diversity in the repertoires of degradosome RNases observed in many bacteria. Indeed, as we recently reported, while 26% of bacteria only possess RNase E (like *E. coli*), 27% have both RNase J and RNase Y (like *H. pylori*) and 19% have both RNase E and RNase J (like *M. tuberculosis*) (1). Thus, it is most probable that different types of degradosomes with different localizations co-exist in numerous bacteria pointing to interesting questions on their respective functions, potential cross-talk and target specificities.

Fluorescent fusions of both partners of the RNA degradosome were analyzed in live *H. pylori* cells by confocal microscopy. RNase J-GFP formed bright foci at the cell periphery that are non-polar. The corresponding strain does not show a growth defect (*rnj* being an essential gene), and therefore the RNase J-GFP foci do not represent misfolded protein aggregates but rather physiologically relevant structures (34). The RNase J foci, a median of 3 per bacterium in exponential phase by confocal microscopy, suggest that RNA degradosome complexes are located at discrete sites at the cell membrane. Under the same conditions, RhpA-CFP also forms foci at the periphery of the cell. Although we favor the hypothesis of “mixed” RNase J-RhpA degradosome foci, this could not be established since colocalization studies were prevented by the fluorescence background throughout the RhpA-CFP expressing cells, that we could attribute to partial membrane delocalization of the RhpA-CFP fusion.

The RNase J-GFP fusion was further used to identify factors that modify its cellular distribution. We identified several factors important for foci formation, interestingly none of them significantly impacted the membrane localization of RNase J and of RhpA. We found that RNase J foci formation was impacted by several mutations (Δ*rhpA*, Δ*rny*, Δ*floA*). The number of RNase J foci per cell was slightly diminished in the absence of its RhpA partner. In the absence of RNase Y, the number of foci per cell raises by 22%. One plausible explanation, supported by the antibiotic treatment data, is that foci formation depends on the amount of free RNA and would, in the mutant deficient in RNase Y, be promoted by the accumulation of its RNA targets. In the absence of the sole flotillin of *H. pylori*, the number of foci per cell was reduced by 46%, showing that this membrane scaffolding protein promotes foci formation. In *S. aureus*, FloA affects the function and oligomeric state of the membrane-bound RNase Y (28). The *H. pylori ΔfloA* mutant presents no growth defect suggesting that RNase J activity is not significantly affected. However, FloA-rich membrane regions might act as scaffolds to bring molecules together maybe fostering oligomerization and foci formation.

We examined how the exposure of *H. pylori* cells to three different antibiotics impacts the formation of RNase J-GFP foci. These treatments do not change the cellular localization of RNase J-GFP. Rifampicin, a transcription inhibitor, and chloramphenicol, a translation inhibitor that locks ribosomes on mRNAs, are known to lower the intracellular concentration of untranslated/free RNAs. These two antibiotics cause a significant reduction in the number of foci per cell (the effect being more pronounced with rifampicin). Puromycin, an antibiotic that leads to the dissociation of the ribosomal subunits, thereby increasing the amounts of untranslated free mRNAs, does not significantly change the number of foci per cell. Altogether, our results show that RNA does not determine the membrane localization of RNase J but rather suggest that free/untranslated RNA is a factor promoting RNase J foci formation. This also shows that not all RNase J proteins present at the membrane are embedded into foci, an aspect that we were able to tackle with super-resolution microscopy.

In *B. subtilis*, rifampicin treatment does not change RNase Y membrane localization, although it modifies the number of RNase Y foci and results in a complete delocalization of RNase J1 and J2 (26, 29). In *E. coli*, rifampicin treatment results in disappearance of RNase E-YFP foci and an increased rate of RNase E membrane diffusion (20). In *C. crescentus*, using 3D single-particle tracking and super-resolution microscopy, it was observed that the number of confined/clustered RNase E molecules decreases upon rifampicin treatment (22). Another study reported that the number of *C. crescentus* RNase E foci was reduced upon rifampicin, chloramphenicol or tetracycline treatment and slightly increased upon puromycin treatment (23). Thus, untranslated RNA appears to play an important role in RNA degradosome foci formation/clustering in distant microorganisms (*H. pylori, E. coli* and *C. crescentus*) and for unrelated RNases (RNase J and RNase E) pointing to evolutionary conserved features.

Our data discussed so far suggest that the formation of RNase J foci is not constitutive, we thus wanted to investigate this under normal physiological conditions, like growth phase. By confocal microscopy, the number of foci was found to diminish during *H. pylori* growth phase reaching almost no detectable foci in stationary phase. Using Western blot, we found that, in stationary phase, the total amount of proteins RNase J-GFP is slightly diminished while its membrane localization is maintained. To have a better insight into the evolution of foci during growth and to better characterize these structures, super-resolution dSTORM, that allows visualization of single molecules, was applied. In agreement with our previous results, the number of RNase J molecules per cell in stationary phase was found to be approximately half that in exponential phase. With the higher sensitivity of dSTORM, we detected more foci than with confocal microscopy, with a median number of 4 foci per cell versus 2, in exponential and stationary phase, respectively. Between exponential and stationary phase, the median volume of the foci is decreased by nine-fold and contains half the number of RNase J molecules. Thus, the foci volume is not linearly proportional to the number of RNase J molecules it contains suggesting that the composition of foci and possibly its activity changes during growth. Importantly, the mean proportion of RNase J molecules that take part of foci remains the same (about 40%) between exponential and stationary phase.

Our dSTORM data allowed us to measure the changes of *H. pylori* cell volume as a function of growth and we found, similarly to what was established in *E. coli*, that the volume is approx. two times smaller in stationary phase. We then calculated that the RNase J “cellular concentration” and number of foci per cell are approx. constant over growth and conclude that the RNase J foci are undergoing a process of cell size scaling. This is particularly interesting since eukaryotic ribonucleoprotein granules have been shown to depend on cell size for assembly (35). Such scaling processes allow the cells to adapt their functions within organelles or membrane-less organelles to their volume changes during growth.

These data, and the fact that the membrane localization of RNase J is not modified along the growth phase while the formation of foci is, suggest that foci assembly is indeed a regulated process and is not due to spontaneous assembly of proteins. Our results also suggest that free RNA contributes to foci formation, through a mechanism that remains to be elucidated. Bacterial mRNA concentration is known to be reduced in stationary phase (36) and could thus constitute one important factor controlling the evolution of foci during growth in *H. pylori*.

A dual pattern of behavior during exponential and stationary phase has been proposed for RNases J1 and J2 from *Streptococcus pyogenes*, and it has been suggested that these RNases might be less active during stationary phase (37). It was also shown that, in *S. pyogenes*, RNase J1/2 are only necessary for the initial step of decay of the transcripts that are degraded in stationary phase (37).

The *E. coli* and *C. crescentus* degradosomes were also found to assemble into foci or clusters (20, 22, 23). However, there are some differences with the degradosome of *H. pylori*. First, the other clusters are highly dynamic while those of *H. pylori* are, under the tested conditions, static. Second, the cytoplasmic clusters of the RNase E-based *C. crescentus* degradosome have LLPS properties similar to eukaryotic messenger ribonucleoprotein (mRNP) granules (like p-bodies or stress granules). Despite their differences, the three types of clustered bacterial degradosomes reported so far (including the one of *H. pylori*) present striking similarities with these eukaryotic structures [some of which being static (38, 39)]; they all form compartmentalized structures within the cell, their formation is regulated and promoted by RNA, and for *H. pylori* they seem to be scaled to the size of the cells. The formation of LLPS structures has been frequently linked to the presence of RNA-binding proteins and proteins with intrinsically disordered regions (IDRs) that can act as intermolecular interaction hotspots. It is known that the C-terminal domain of *Eco*RNase E is an IDR (40). Using the IUPred2A algorithm to predict IDRs (41), such a region was predicted in the N-terminal part of *H. pylori’*s RNase J (100 first aa, Fig. S1C), a region that is specific to *Helicobacter* RNase J proteins. An IDR is also predicted in the C-terminus (last 60 aa) of RhpA (Fig. S1D). These predictions remain to be experimentally validated, however this might be complicated by the fact that, as we previously showed, the N-terminal region of RNase J is important for its activity (10, 11). Future experiments will be required to explore whether these regions play a role in the formation of the RNA degradosome foci in *H. pylori*.

It is still not clear whether the bacterial degradosome foci correspond to active RNA degradation hubs or a storage form of the enzymes and/or RNA molecules or other proteins. Surprisingly, after many years, this question is still partially open for p-bodies and stress granules. Recent data of *C. crescentus* however suggest that they are active structures (24).

We propose a working model for the dual localization of the degradosome (Fig. 7). The membrane association of the degradosome could be a manner to compartmentalize RNA degradation and an active form of the degradosome would be clustered into foci. This could allow post-transcriptional regulation of a subcategory of genes whose mRNAs are directed to the foci by an unknown mechanism while also providing a spatio-temporal delay to allow transcribed mRNAs to be translated before their degradation. The ribosome-associated degradosomes (be they at the membrane and forming foci or not) could be in charge of degrading RNA molecules that need to be tightly regulated and would be deleterious if left unchecked, they could degrade defective RNA molecules or play a role in ribosomal RNA maturation, as has been shown for RhpA (12).

**Figure 7.**
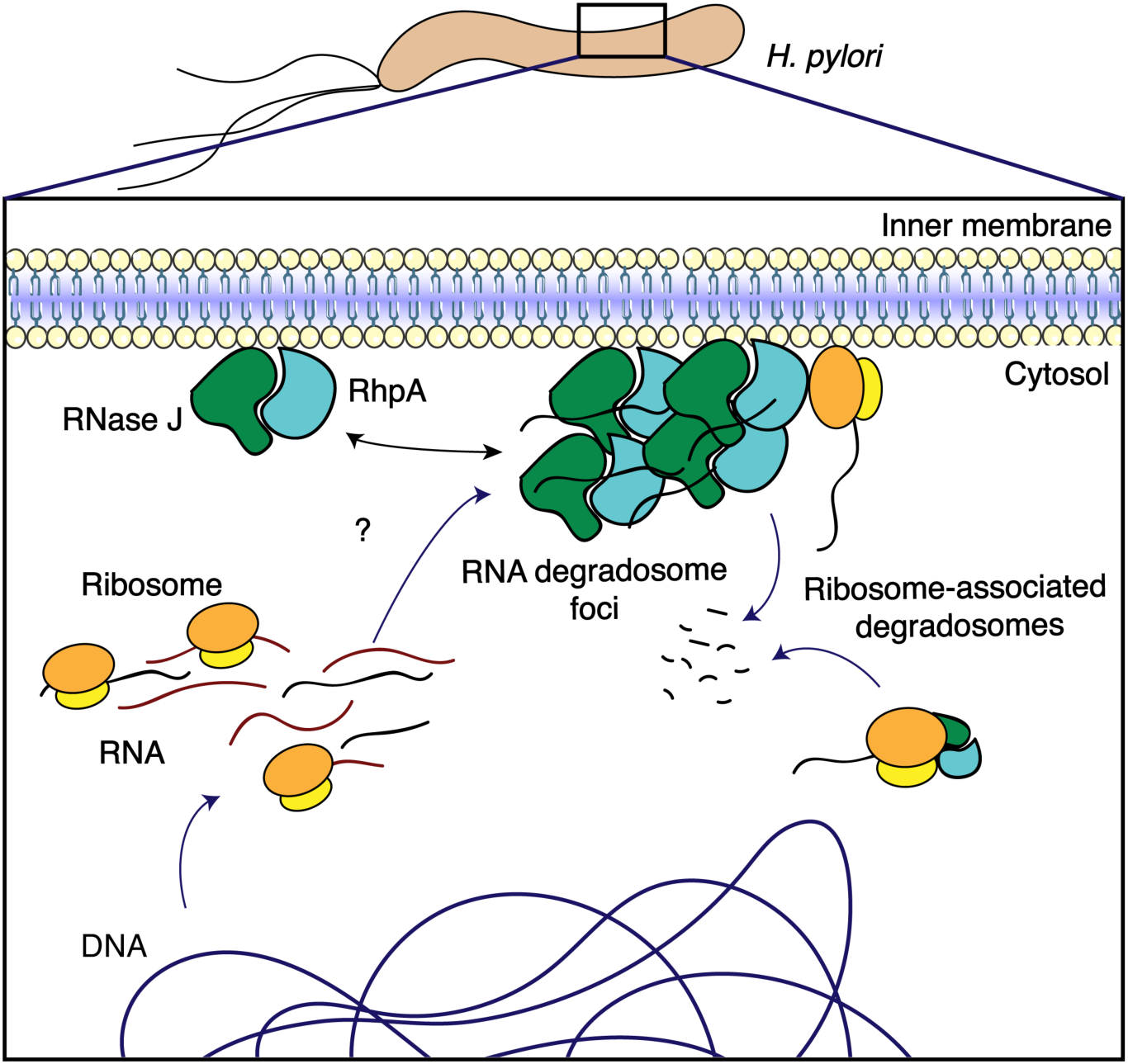
Model for the regulation of the RNA degradosome foci in *H. pylori*. RNase J and RhpA, the two protein components of the RNA degradosome are associated with the *H. pylori* inner membrane. A minor proportion of these proteins is associated to translating ribosomes, which could be either cytoplasmic or at the membrane. RNase J and RhpA assemble into foci at the membrane, probably together. Based on our data, we propose a model where foci represent the active form of the RNA degradosome, constituting RNA degradation hubs. Comparatively, the complexes outside foci would retain little or no activity. In this model, target RNA molecules would be directed to the foci for degradation by an unknown mechanism, providing in addition a spatio-temporal delay that allows for their translation before their degradation. The RNA degradosomes associated with ribosomes could either be involved in rRNA maturation or in coupling between translation and mRNA degradation.

In conclusion, we have discovered and characterized the membrane localization and clustering of the RNA degradosome in the important pathogen *H. pylori*. We propose that the balance between this localization, its clustering and ribosomal association are major control levels of its activity and specificity that might be more generally relevant and open many exciting perspectives of research by analogy with the equivalent eukaryotic structures.

## Materials and methods

### Bacterial strains and growth conditions

The *H. pylori* strains used in this study (Table S1) are derivatives from B128 (42, 43). Plasmids (Table S1) used to create mutants of *H. pylori* were constructed and amplified using *Escherichia coli* one-shot top10 or DH5α strains (Thermofisher). *H. pylori* strains were grown on Blood Agar Base 2 (Oxoid) plates supplemented with 10% defibrinated horse blood and with the following antibiotics-antifungal cocktail: amphotericin B 2.5 μg.ml^-1^, polymyxin B 0.31 μg.ml^-1^, trimethoprim 6.25 μg.ml^-1^ and vancomycin 12.5 μg.ml^-1^. Selection of *H. pylori* mutants was performed using kanamycin 20 μg.ml^-1^, apramycin 10μg.ml^-1^ or chloramphenicol 10 μg.ml^-1^. For liquid cultures, we used Brucella broth supplemented with 10% Fetal Calf Serum (FCS) (Eurobio), the antibiotics-antifungal cocktail and the selective antibiotic when necessary. *H. pylori* cells were grown at 37°C under microaerophilic atmosphere conditions (6% O_2_, 10% CO_2_, 84% N_2_) using an Anoxomat (MART Microbiology) atmosphere generator.

### Molecular techniques

Molecular biology experiments were performed according to standard procedures (44) and the supplier (Fermentas) recommendations. NucleoBond Xtra Midi Kit (Macherey-Nagel) and QIAamp DNA Mini Kit (Qiagen) were used for plasmid preparations and *H. pylori* genomic DNA extractions, respectively. PCR were performed either with DreamTaq DNA polymerase (Thermofisher), or with Q5 High fidelity DNA polymerase (NEB) when the product required high fidelity polymerase.

### Construction of H. pylori mutants and fusions

Chromosomal deletion of the entire genes encoding RNase Y (HPB128_186g30), RhpA (HPB128_21g22) and flotillin (HPB128_21g23) was performed in strain B128. Briefly, fragments of about 500 bp upstream and downstream of the target gene were amplified by PCR (oligonucleotides are listed in Table S1) and spliced into a non-polar kanamycin, chloramphenicol or apramycin resistance cassette by using the Gibson isothermal assembly kit (NEB) followed by PCR amplification using the primers from the extremities. All *H. pylori* mutants were obtained by natural transformation [as described previously (45)] with the PCR fragments obtained as above. Selection of chromosomal allelic exchange resulting in gene deletion was performed with the corresponding antibiotic. Deletion of the genes of interest and correct insertion of cassettes were verified by PCR and sequencing of the gene region. RNase J-GFP, RNase J-FLAG, RhpA-CFP and L9-V5 fusions were also constructed using the isothermal assembly technique(46). Briefly, target genes, GFPmut2 (47) or V5 tag (48) followed by the kanamycin resistance cassette; SCFP (49) followed by the chloramphenicol resistance cassette; or FLAG tag (50) followed by the apramycin resistance cassette, were fused in this order after amplification using complementary primers (described in Fig. S2A). A linker sequence (4 aa, sequence Phe-His-Gly-Ser) was introduced between RNase J/RhpA and the fluorescent proteins in RNase J-GFP and RhpA-CFP fusions. Then, the final construction was amplified by PCR and directly introduced into *H. pylori* by natural transformation. Correct insertion of the fusion and cassette was verified by PCR, sequencing, Western blot and, when relevant, microscopy. Additionally, the *gfpmut2* (51) and *scfp* genes were cloned into the pILL2157 shuttle vector that replicates in *H. pylori* (52) between the *Nde*I and *BamH*I restriction sites under the control of an IPTG inducible promoter. These constructs were transformed into *H. pylori*, verified by PCR and sequencing and served as controls for microscopy analysis.

### Confocal fluorescence microscopy

Bacterial samples taken from cultures with different OD_600nm_ were concentrated or diluted to an OD_600nm_ of 2 and, when necessary, stained. The cultures were labeled with NucBlue® Live Cell Stain Ready Probes™ Reagent (Hoechst 33342, Molecular Probes) for DNA as per the recommendations of the supplier and with 4 µg/mL FM4-64 (Thermo Fisher Scientific) for membranes. For strains carrying the RhpA-CFP fusion, only FM4-64 was used. After 20 minutes of incubation in the dark, bacterial samples were spotted on microscope slides with a thin layer of agarose (1% in PBS). The agarose pad was covered by a coverslip and samples were directly observed at room temperature. Images of strains expressing RNase J-GFPmut2, RhpA-SCFP from the chromosome or GFPmut2 and SCFP from pILL2157 were obtained using a Leica confocal SP8 inverted microscope through an objective 63x NA 1.4 oil immersion.

The detection has been performed by a PMT detector for the NucBlue® channel and HyD for the other fluorophores. Images were analyzed using Fiji (53). Pictures were deconvolved using the classic maximum likelihood algorithm of the Huygens software (SVI, Laapersveld, The Netherlands).

The RNase J-FLAG and RhpA-FLAG expressing strains were fixed and permeabilized with 100% methanol at −20°C for 5 min. Subsequently, the bacteria were washed with PBS and deposited on poly-L-lysine-covered coverslips. The samples were quenched for 30 min at room temperature with 100 mM glycine, and were then blocked with 5% BSA in PBS for 1h at room temperature. They were then incubated with α-FLAG antibody (1:200, Sigma) in 2% BSA in PBS, washed three times with 2% BSA in PBS, incubated with α-mice-AF488 secondary antibody (1:200, Bethyl), washed six times with 2% BSA in PBS and mounted on Fluoromount-G (Thermo Fisher Scientific). No signal was seen when the protocol was carried out on a strain lacking a FLAG tag. Imaging and analysis were performed as described above.

### Foci quantification

To quantify the RNase J-GFP foci in *H. pylori*, images of cells were first segmented manually using the nucleoid signal, which was homogeneous. Only those cells that could be segmented with certainty were taken. RNase J-GFP foci were detected using the blob detector function with scale space (variable target blob size) from scikit-image (54). The particle size range was set to from 1 to 2 pixels (corresponding to about 60-120 nm); the sensitivity of the blob detector was set to 0.006. This sensitivity was kept constant during analysis of all images and adjusted only in a few noisy images to suppress apparent false positives from the background.

To correct for the lateral shift (in the xy-plane) due to chromatic aberrations, we used a standard approach utilizing fiducial markers. Multicolor fluorescent beads (TetraSpeck 0.2 µm microspheres, Thermofisher) were dispersed in buffer, immobilized on a glass coverslip and imaged in the DAPI and GFP channels. Beads were detected in the images and their xy positions were found with sub-pixel resolution using the FIJI plugin TrackMate (55) and for each bead the lateral shift (localization discrepancy between the channels) was calculated; the average shift per bead gives the single shift vector between the GFP and DAPI channels. This vector is used in the following analysis scripts to obtain corrected degradosome positions (GFP channel) relative to the nucleoid positions (DAPI channels). Next, for each RNase J focus, we searched for its nearest nucleoid within a fixed search radius (twice the average nucleoid size). The distance between a nucleoid and a RNase J focus was calculated from RNase J focus position (its center) and a nucleoid centroid. This step typically yields several nucleoid candidates, only one of which is selected, the one whose edge comes the closest to the foci. This allows to determine the number of degradosome foci per nucleoid.

The position of the RNase J foci with respect to the bacterial poles was determined as follows. We defined the medial axis of the nucleoid (spine) using skeletonization function in Python library skimage, and then assigned the values 0 and 1 to either end of the spine (poles). For each RNase J focus, the closest point on the spine of its host nucleoid was found. The length of the spine between this point and the closest pole was divided by the total length of the spine to find the relative position of the RNase J focus along the spine (RP). In this way, if for a RNase J focus RP = 0 or 1, then it is located at one of the poles, whereas if RP = 0.5, then it is located at the middle of the spine. Lastly, the frequencies of foci with RP = 0 or 1 were normalized by the perimeter of the pole tip to compensate for the accumulation of foci due to their rotational freedom around the tip. The code data are available at the following repository https://gitlab.pasteur.fr/iah-public/hpyloridegradosomeanalysis.

### Foci mobility

To quantitatively compare the mobility of the foci in live and fixed cells we used the mean square displacement (MSD) as metrics. For this, foci were first tracked with sub-pixel resolution using the TrackMate plugin in FIJI (55). Resulting tracks were analyzed using MSDanalyzer (56) in Matlab. The trajectories were corrected for drift using velocity correlation in MSDanalyzer.

### Fractionation of *H. pylori*

The cellular fractionation protocol was adapted from (30). *H. pylori* cells were grown to an appropriate OD_600 nm_ (0.7-1.5 for exponential phase or >3 for stationary phase) and then harvested by centrifugation and washed with PBS prior to being resuspended (normalizing to an OD_600 nm_ of 5). Then, bacteria were disrupted by sonication in a lysis buffer containing 10 mM Tris-HCl pH 7.4 (buffer A) and Complete Protease Inhibitor Cocktail (Roche). Cell debris was removed by centrifugation at 16,000 g at 4°C during 10 minutes and supernatants were collected as total extracts. The supernatants were transferred to ultracentrifugation tubes (Polyallomer, Beckman Coulter) of 1.5 mL and then centrifuged 45 min at 100,000g at 4°C in a TLA-100 ultracentrifuge (Beckman Coulter). The supernatant contains the soluble fraction and the pellet corresponds to the total membranes. The pellet was washed once with buffer A and then resuspended in 10 mM Tris-HCl pH 7.5 + 0.1% N-lauroyl-sarcosin (Sigma-Aldrich) and Complete Protease Inhibitor Complete (buffer B). After another ultracentrifugation under the same conditions, the supernatant contains the inner membrane and the pellet the outer membrane, which was resuspended in 10 mM Tris-HCl pH 7.5 + 1% N-lauroyl-sarcosin and Complete Protease Inhibitor Complete (buffer C). At each step, each sample is suspended in the same volume to obtain samples from the same number of bacteria. To determine the nature of the interaction of RhpA and RNase J with the inner membrane of *H. pylori*, we tested different treatments as described in (57). Peripheral membrane proteins dissociate from the membrane when treated with a polar reagent that does not disrupt the lipid bilayer (urea 6 M), exposed to extreme alkaline pH (Na_2_CO_3_ 100 mM pH 11) or to high salt concentration (NaCl 2 M). After ultracentrifugation for 45 min at 100,000 g, peripheral proteins are found in the supernatant while integral membrane proteins are found in the pellet. Finally, treatments with 1 µg/ml RNase A (ThermoFisher) or 20 mM EDTA (Sigma-Aldrich) were performed by adding them to the cells just after sonication and were maintained during the first ultracentrifugation.

### Antibiotic treatments

To examine the effect of antibiotics on the subcellular localization of RNase J and RhpA and the RNase J-GFP foci formation, rifampicin, chloramphenicol or puromycin were added at 100 µg/mL, 200 µg/mL and 150 µg/mL respectively to cultures in late exponential phase, as in (23). Cultures were further incubated during 30 min under microaerobic atmosphere at 37°C under agitation. The cultures were then collected and treated for microscopy and/or fractionation.

### Western blotting

Proteins were loaded and separated on a 4-20% Mini-Protean TGX Stain-Free precast protein gel (BioRad) and subsequently electrotransferred to a polyvinylidene difluoride (PVDF) membrane (Biorad) with the TransBlot Turbo system (Biorad). The *H. pylori* RhpA, RNase J, AmiE, MotB and BabA proteins were detected with rabbit polyclonal antibodies α-RhpA, α-RNase J (10), α-AmiE (58), α-MotB (gift of N. Buddelmeijer) and α-BabA (59) at the respective dilutions of 1:5,000, 1:500, 1:500, 1:500 and 1:10,000. Goat anti-rabbit IgG-HRP (Santa Cruz) was used as secondary antibody at 1:10,000 dilution and the detection was achieved with the ECL Femto reagent (Thermo Fisher). The V5 tag was detected with an anti-V5 antibody coupled with HRP (SantaCruz) (1:5,000).

### Sample preparation for fluorescence nanoscopy using single-molecule localization microscopy

Bacterial suspensions of the RNase J-GFP expressing strain in exponential growth phase (after 18 hours of culture) or in stationary phase (after 40h of culture), were fixed in 1% paraformaldehyde (Sigma) in PBS for 5 min at room temperature. Following three washes in PBS, bacteria were permeabilized using 0.05% Triton X-100 in PBS. Anti-GFP nanobodies coupled with Cy5 (Fluotag®-Q, NanoTag Biotechnologies GmbH) were then added (1:250) to the bacterial suspensions and incubated for 1h at 37°C. The cells were then washed and treated with 2 µg/mL WGA-AF555 (Thermo Fisher Scientific) in PBS. In order to (i) immobilize the bacterial cells onto the coverslips (#1.5H, Marienfeld, Germany) and (ii) induce the photoswitching of Cy5, the bacteria were seeded onto STORM buffer-based pads (Blinking Pad Kit, Abbelight, Paris, France).

### Single-molecule localization microscopy imaging and analysis

2D and 3D images were taken using an inverted bright-field Olympus IX83 microscope equipped with a 100x oil-immersion objective with a high numerical aperture (1.49). To perform fluorescence nanoscopy experiments, the SAFe360 module (Abbelight, France) was added to the camera port of the microscope. This detection module couples Single-Molecule Localization Microscopy (SMLM), Supercritical Angle Fluorescence (SAF) and astigmatism (60, 61) in a dual-view setup coupled with sCMOS cameras (Orcaflash v4, Hamamatsu). Prior to each acquisition, bright-light and diffraction-limited images were acquired. Using continuous excitation at 639 nm in the HiLo mode, most of the Cy5 molecules were induced into a dark state until a sufficient density was obtained (typically 1 to 5 molecules per bacterium per frame). Image series (5000 frames) were recorded with a 50-ms exposure time. Raw images and resulting coordinate tables were processed and analyzed using NEO SAFe software (Abbelight, France). Analysis of high-density regions of localizations was performed using the density-based clustering algorithm DBSCAN (62), using a distance parameter of 75 nm and a minimum number of points parameter of 5.

The volume of *H. pylori* cells was calculated with the d-STORM images and based on the XYZ coordinates of the detections of the membrane marker (WGA-AF555). Clusters of detections belonging to a bacterium were identified with the XY coordinates and the clusters were refined by filtering by the number of points in a cluster and the radius of gyration. Each cluster was converted into a 2D polygon using the Python library alphashape and the 2D area (A) of the polygon was calculated from the XY-coordinates of the underlying cluster. Using this area, the volume (V) was calculated using the expression as V=H*A, where H is the height of the bacteria. The height was estimated from the span of coordinates in the Z dimension of the clusters (600 nm). The volume of the bacteria might be slightly overestimated due to the localization errors in imaging, however this should not influence the ratio of exponential to stationary growing cells.

### Statistical analysis

To compare the numbers of degradosome foci per cell between the different conditions, we used the non-parametric Mood Median test and p-values smaller than 0.05 were considered as significant. The differences between the median values derived from super-resolution microscopy were analyzed with the Mann-Whitney test and p-values smaller than 0.05 were considered as significant.

## Acknowledgements

ATA is part of the Pasteur - Paris University (PPU) International PhD Program. This project has received funding from the European Union’s Horizon 2020 research and innovation programme under the Marie Sklodowska-Curie grant agreement Nº 665807, and from the Institut Carnot Pasteur Microbes & Santé. EG was funded by a Université Paris Diderot Paris 7 fellowship. Funding was provided by the Agence Nationale de la Recherche [ANR 09 BLAN 0287 01, PyloRNA to H.D.R.] and the [ANR-12-BSV5-0025-02 to HDR and LEM]. We thank the Institut Pasteur for the Bourse Roux research fellowships (to LEM) and Janssen for financial support. Support was provided by “Fondation pour la Recherche Médicale” for the grant DBF20161136767 to HDR and the Pasteur-Weizmann Consortium of “The Roles of Noncoding RNAs in Regulation of Microbial Life Styles and Virulence” to HDR. We are also thankful to Marie Pastemps and Frédéric Fischer for their advice and help in the experiments, and to Julien Fernandes for his help with microscopy images. We also thank Ivo G. Boneca and Wolfgang Fischer for gift of antibodies, and Nienke Buddelmeijer for the gift of CFP and antibodies. The UtechS Photonic BioImaging (Imagopole), C2RT, Institut Pasteur was supported by the French National Research Agency (France BioImaging; ANR-10–INSB–04; Investments for the Future). We thank the company Abbelight and in particular Rym Boudjemaa for super-resolution image acquisition, data processing and analysis. Some figures were generated with images from Servier Medical Art (https://smart.servier.com/), licensed under the Creative Commons Attribution 3.0 Unported License (http://creativecommons.org/license/by/3.0/).

## Author contributions

Conceptualization: ATA EG DE HDR

Methodology: ATA EG ET LEM DE HDR

Software: DE

Validation: ATA EG

Formal analysis: ATA EG DE HDR

Investigation: ATA EG LEM ET

Resources: DE HDR

Data curation: DE

Writing ± original draft: ATA EG DE HDR

Writing ± review & editing: ATA EG ET LEM DE HDR

Visualization: ATA EG

Supervision: HDR

Project administration: HDR

Funding Acquisition: HDR

## References

1. Tejada-Arranz A, de Crécy-Lagard V, de Reuse H. 2020. Bacterial RNA Degradosomes: Molecular Machines under Tight Control. Trends Biochem Sci 45:42–57.

2. Bandyra KJ, Bouvier M, Carpousis AJ, Luisi BF. 2013. The social fabric of the RNA degradosome. Biochim Biophys Acta 1829:514–22.

3. Vanzo NF, Li YS, Py B, Blum E, Higgins CF, Raynal LC, Krisch HM, Carpousis AJ. 1998. Ribonuclease E organizes the protein interactions in the Escherichia coli RNA degradosome. Genes Dev 12:2770–81.

4. Hardwick SW, Chan VSY, Broadhurst RW, Luisi BF. 2011. An RNA degradosome assembly in Caulobacter crescentus. Nucleic Acids Res 39:1449–59.

5. Plocinski P, Macios M, Houghton J, Niemiec E, Plocinska R, Brzostek A, Słomka M, Dziadek J, Young D, Dziembowski A. 2019. Proteomic and transcriptomic experiments reveal an essential role of RNA degradosome complexes in shaping the transcriptome of Mycobacterium tuberculosis. Nucleic Acids Res 47:5892–5905.

6. Carpousis AJ. 2007. The RNA Degradosome of *Escherichia coli* : An mRNA-Degrading Machine Assembled on RNase E. Annu Rev Microbiol 61:71–87.

7. Mathy N, Bénard L, Pellegrini O, Daou R, Wen T, Condon C. 2007. 5′-to-3′ Exoribonuclease Activity in Bacteria: Role of RNase J1 in rRNA Maturation and 5′ Stability of mRNA. Cell 129:681–692.

8. Shahbabian K, Jamalli A, Zig L, Putzer H. 2009. RNase Y, a novel endoribonuclease, initiates riboswitch turnover in Bacillus subtilis. EMBO J 28:3523–3533.

9. Durand S, Condon C. 2018. RNases and Helicases in Gram-Positive Bacteria. Microbiol Spectr 6:37–53.

10. Redko Y, Aubert S, Stachowicz A, Lenormand P, Namane A, Darfeuille F, Thibonnier M, De Reuse H. 2013. A minimal bacterial RNase J-based degradosome is associated with translating ribosomes. Nucleic Acids Res 41:288–301.

11. Redko Y, Galtier E, Arnion H, Darfeuille F, Sismeiro O, Coppée J-Y, Médigue C, Weiman M, Cruveiller S, De Reuse H. 2016. RNase J depletion leads to massive changes in mRNA abundance in *Helicobacter pylori*. RNA Biol 13:243–253.

12. El Mortaji L, Aubert S, Galtier E, Schmitt C, Anger K, Redko Y, Quentin Y, De Reuse H. 2018. The sole DEAD-box RNA helicase of the gastric pathogen Helicobacter pylori is essential for colonization. MBio 9:e02071–17.

13. Salama NR, Hartung ML, Müller A. 2013. Life in the human stomach: persistence strategies of the bacterial pathogen Helicobacter pylori. Nat Rev Microbiol 11:385–99.

14. Amieva M, Peek RM. 2016. Pathobiology of Helicobacter pylori–Induced Gastric Cancer. Gastroenterology 150:64–78.

15. Sharma CM, Hoffmann S, Darfeuille F, Reignier J, Findeiß S, Sittka A, Chabas S, Reiche K, Hackermüller J, Reinhardt R, Stadler PF, Vogel J. 2010. The primary transcriptome of the major human pathogen Helicobacter pylori. Nature 464:250–255.

16. Khemici V, Poljak L, Luisi BF, Carpousis AJ. 2008. The RNase E of *Escherichia coli* is a membrane-binding protein. Mol Microbiol 70:799–813.

17. Liou GG, Jane WN, Cohen SN, Lin NS, Lin-Chao S. 2001. RNA degradosomes exist in vivo in Escherichia coli as multicomponent complexes associated with the cytoplasmic membrane via the N-terminal region of ribonuclease E. Proc Natl Acad Sci U S A 98:63–8.

18. Hadjeras L, Poljak L, Bouvier M, Morin-Ogier Q, Canal I, Cocaign-Bousquet M, Girbal L, Carpousis AJ. 2019. Detachment of the RNA degradosome from the inner membrane of *Escherichia coli* results in a global slowdown of mRNA degradation, proteolysis of RN ase E and increased turnover of ribosome-free transcripts. Mol Microbiol mmi. 14248.

19. Moffitt JR, Pandey S, Boettiger AN, Wang S, Zhuang X. 2016. Spatial organization shapes the turnover of a bacterial transcriptome. Elife 5.

20. Strahl H, Turlan C, Khalid S, Bond PJ, Kebalo J-M, Peyron P, Poljak L, Bouvier M, Hamoen L, Luisi BF, Carpousis AJ. 2015. Membrane recognition and dynamics of the RNA degradosome. PLoS Genet 11:e1004961.

21. Tsai Y-C, Du D, Domínguez-Malfavón L, Dimastrogiovanni D, Cross J, Callaghan AJ, García-Mena J, Luisi BF. 2012. Recognition of the 70S ribosome and polysome by the RNA degradosome in Escherichia coli. Nucleic Acids Res 40:10417–10431.

22. Bayas CA, Wang J, Lee MK, Schrader JM, Shapiro L, Moerner WE. 2018. Spatial organization and dynamics of RNase E and ribosomes in Caulobacter crescentus. Proc Natl Acad Sci U S A 115:E3712–E3721.

23. Al-Husini N, Tomares DT, Bitar O, Childers WS, Schrader JM. 2018. α-Proteobacterial RNA Degradosomes Assemble Liquid-Liquid Phase-Separated RNP Bodies. Mol Cell 71:1027–1039.e14.

24. Al-Husini N, Tomares DT, Pfaffenberger ZJ, Muthunayake NS, Samad MA, Zuo T, Bitar O, Aretakis JR, Bharmal M-HM, Gega A, Biteen JS, Childers WS, Schrader JM. 2020. BR-Bodies Provide Selectively Permeable Condensates that Stimulate mRNA Decay and Prevent Release of Decay Intermediates. Mol Cell 78:1–13.

25. Hunt A, Rawlins JP, Thomaides HB, Errington J. 2006. Functional analysis of 11 putative essential genes in Bacillus subtilis. Microbiology 152:2895–2907.

26. Cascante-Estepa N, Gunka K, Stülke J. 2016. Localization of Components of the RNA-Degrading Machine in Bacillus subtilis. Front Microbiol 07:1492.

27. Khemici V, Prados J, Linder P, Redder P. 2015. Decay-Initiating Endoribonucleolytic Cleavage by RNase Y Is Kept under Tight Control via Sequence Preference and Sub-cellular Localisation. PLOS Genet 11:e1005577.

28. Koch G, Wermser C, Acosta IC, Kricks L, Stengel ST, Yepes A, Lopez D. 2017. Attenuating Staphylococcus aureus Virulence by Targeting Flotillin Protein Scaffold Activity. Cell Chem Biol 24:845–857.e6.

29. Hamouche L, Billaudeau C, Rocca A, Chastanet A, Ngo S, Laalami S, Putzer H. 2020. Dynamic Membrane Localization of RNase Y in Bacillus subtilis. MBio 11.

30. Kutter S, Buhrdorf R, Haas J, Schneider-Brachert W, Haas R, Fischer W. 2008. Protein subassemblies of the Helicobacter pylori Cag type IV secretion system revealed by localization and interaction studies. J Bacteriol 190:2161–71.

31. Hutton ML, D’Costa K, Rossiter AE, Wang L, Turner L, Steer DL, Masters SL, Croker BA, Kaparakis-Liaskos M, Ferrero RL. 2017. A Helicobacter pylori Homolog of Eukaryotic Flotillin Is Involved in Cholesterol Accumulation, Epithelial Cell Responses and Host Colonization. Front Cell Infect Microbiol 7:219.

32. Šiková M, Wiedermannová J, Převorovský M, Barvík I, Sudzinová P, Kofronová O, Benada O, Šanderová H, Condon C, Krásný L. 2020. The torpedo effect in Bacillus subtilis: RNase J1 resolves stalled transcription complexes. EMBO J 39:e102500.

33. Mu R, Shinde P, Zou Z, Kreth J, Merritt J. 2019. Examining the Protein Interactome and Subcellular Localization of RNase J2 Complexes in Streptococcus mutans. Front Microbiol 10:2150.

34. Zietkiewicz S, Liberek K. 2010. Dispose to the pole-protein aggregation control in bacteria. EMBO J 29:869–70.

35. Brangwynne CP. 2013. Phase transitions and size scaling of membrane-less organelles. J Cell Biol 203:875–881.

36. Chen H, Shiroguchi K, Ge H, Xie XS. 2015. Genome-wide study of mRNA degradation and transcript elongation in E scherichia coli. Mol Syst Biol 11:781–781.

37. Bugrysheva J V., Scott JR. 2010. The ribonucleases J1 and J2 are essential for growth and have independent roles in mRNA decay in *Streptococcus pyogenes*. Mol Microbiol 75:731–743.

38. Kroschwald S, Maharana S, Mateju D, Malinovska L, Nüske E, Poser I, Richter D, Alberti S. 2015. Promiscuous interactions and protein disaggregases determine the material state of stress-inducible RNP granules. Elife 4:e06807.

39. Kroschwald S, Munder MC, Maharana S, Franzmann TM, Richter D, Ruer M, Hyman AA, Alberti S. 2018. Different Material States of Pub1 Condensates Define Distinct Modes of Stress Adaptation and Recovery. Cell Rep 23:3327–3339.

40. Aït-Bara S, Carpousis AJ, Quentin Y. 2015. RNase E in the γ-Proteobacteria: conservation of intrinsically disordered noncatalytic region and molecular evolution of microdomains. Mol Genet Genomics 290:847–862.

41. Mészáros B, Erdos G, Dosztányi Z. 2018. IUPred2A: Context-Dependent Prediction of Protein Disorder as a Function of Redox State and Protein Binding. Nucleic Acids Res 46.

42. McClain MS, Shaffer CL, Israel DA, Peek RM, Cover TL. 2009. Genome sequence analysis of Helicobacter pylori strains associated with gastric ulceration and gastric cancer. BMC Genomics 10:3.

43. Farnbacher M, Jahns T, Willrodt D, Daniel R, Haas R, Goesmann A, Kurtz S, Rieder G. 2010. Sequencing, annotation, and comparative genome analysis of the gerbil-adapted Helicobacter pylori strain B8. BMC Genomics 11:335.

44. Sambrook J, Russell DW (David W. 2001. Molecular cloning : a laboratory manual. Cold Spring Harbor Laboratory Press.

45. Bury-Mone S, Skouloubris S, Dauga C, Thiberge J-M, Dailidiene D, Berg DE, Labigne A, De Reuse H. 2003. Presence of Active Aliphatic Amidases in Helicobacter Species Able To Colonize the Stomach. Infect Immun 71:5613–5622.

46. Gibson DG, Young L, Chuang R-Y, Venter JC, Hutchison CA, Smith HO. 2009. Enzymatic assembly of DNA molecules up to several hundred kilobases. Nat Methods 6:343–345.

47. Cormack BP, Valdivia RH, Falkow S. 1996. FACS-optimized mutants of the green fluorescent protein (GFP). Gene 173:33–38.

48. Hanke T, Young DF, Doyle C, Jones I, Randall RE. 1995. Attachment of an oligopeptide epitope to the C-terminus of recombinant SIV gp160 facilitates the construction of SMAA complexes while preserving CD4 binding. J Virol Methods 53:149–56.

49. Kremers G-J, Goedhart J, van Munster EB, Gadella TWJ. 2006. Cyan and Yellow Super Fluorescent Proteins with Improved Brightness, Protein Folding, and FRET Förster Radius. Biochemistry 45:6570–6580.

50. Hopp TP, Prickett KS, Price VL, Libby RT, March CJ, Pat Cerretti D, Urdal DL, Conlon PJ. 1988. A Short Polypeptide Marker Sequence Useful for Recombinant Protein Identification and Purification. Bio/Technology 6:1204–1210.

51. Babu M, Butland G, Pogoutse O, Li J, Greenblatt JF, Emili A. 2009. Sequential peptide affinity purification system for the systematic isolation and identification of protein complexes from Escherichia coli. Methods Mol Biol 564:373–400.

52. Boneca IG, Ecobichon C, Chaput C, Mathieu A, Guadagnini S, Prévost M-C, Colland F, Labigne A, de Reuse H. 2008. Development of inducible systems to engineer conditional mutants of essential genes of Helicobacter pylori. Appl Environ Microbiol 74:2095–102.

53. Schindelin J, Arganda-Carreras I, Frise E, Kaynig V, Longair M, Pietzsch T, Preibisch S, Rueden C, Saalfeld S, Schmid B, Tinevez J-Y, White DJ, Hartenstein V, Eliceiri K, Tomancak P, Cardona A. 2012. Fiji: an open-source platform for biological-image analysis. Nat Methods 9:676–682.

54. van der Walt S, Schönberger JL, Nunez-Iglesias J, Boulogne F, Warner JD, Yager N, Gouillart E, Yu T. 2014. scikit-image: image processing in Python. PeerJ 2:e453.

55. Tinevez J-Y, Perry N, Schindelin J, Hoopes GM, Reynolds GD, Laplantine E, Bednarek SY, Shorte SL, Eliceiri KW. 2017. TrackMate: An open and extensible platform for single-particle tracking. Methods 115:80–90.

56. Tarantino N, Tinevez J-Y, Crowell EF, Boisson B, Henriques R, Mhlanga M, Agou F, Israël A, Laplantine E. 2014. TNF and IL-1 exhibit distinct ubiquitin requirements for inducing NEMO–IKK supramolecular structures. J Cell Biol 204:231–245.

57. Papanastasiou M, Orfanoudaki G, Koukaki M, Kountourakis N, Sardis MF, Aivaliotis M, Karamanou S, Economou A. 2013. The *Escherichia coli* Peripheral Inner Membrane Proteome. Mol Cell Proteomics 12:599–610.

58. Skouloubris S, Labigne A, De Reuse H. 2001. The AmiE aliphatic amidase and AmiF formamidase of Helicobacter pylori: natural evolution of two enzyme paralogues. Mol Microbiol 40:596–609.

59. Odenbreit S, Kavermann H, Püls J, Haas R. 2002. CagA tyrosine phosphorylation and interleukin-8 induction by Helicobacter pylori are independent from AlpAB, HopZ and Bab group outer membrane proteins. Int J Med Microbiol 292:257–266.

60. Bourg N, Mayet C, Dupuis G, Barroca T, Bon P, Lécart S, Fort E, Lévêque-Fort S. 2015. Direct optical nanoscopy with axially localized detection. Nat Photonics 9:587–593.

61. Cabriel C, Bourg N, Jouchet P, Dupuis G, Leterrier C, Baron A, Badet-Denisot M-A, Vauzeilles B, Fort E, Lévêque-Fort S. 2019. Combining 3D single molecule localization strategies for reproducible bioimaging. Nat Commun 10:1980.

62. Ester M, Kriegel H-P, Sander J, Xu X. 1996. A Density-Based Algorithm for Discovering Clusters in Large Spatial Databases with Noise.

