## Supplementary Figures for "The RNase J-based RNA degradosome is compartmentalized in the gastric pathogen *Helicobacter pylori*"

**Figure S1**

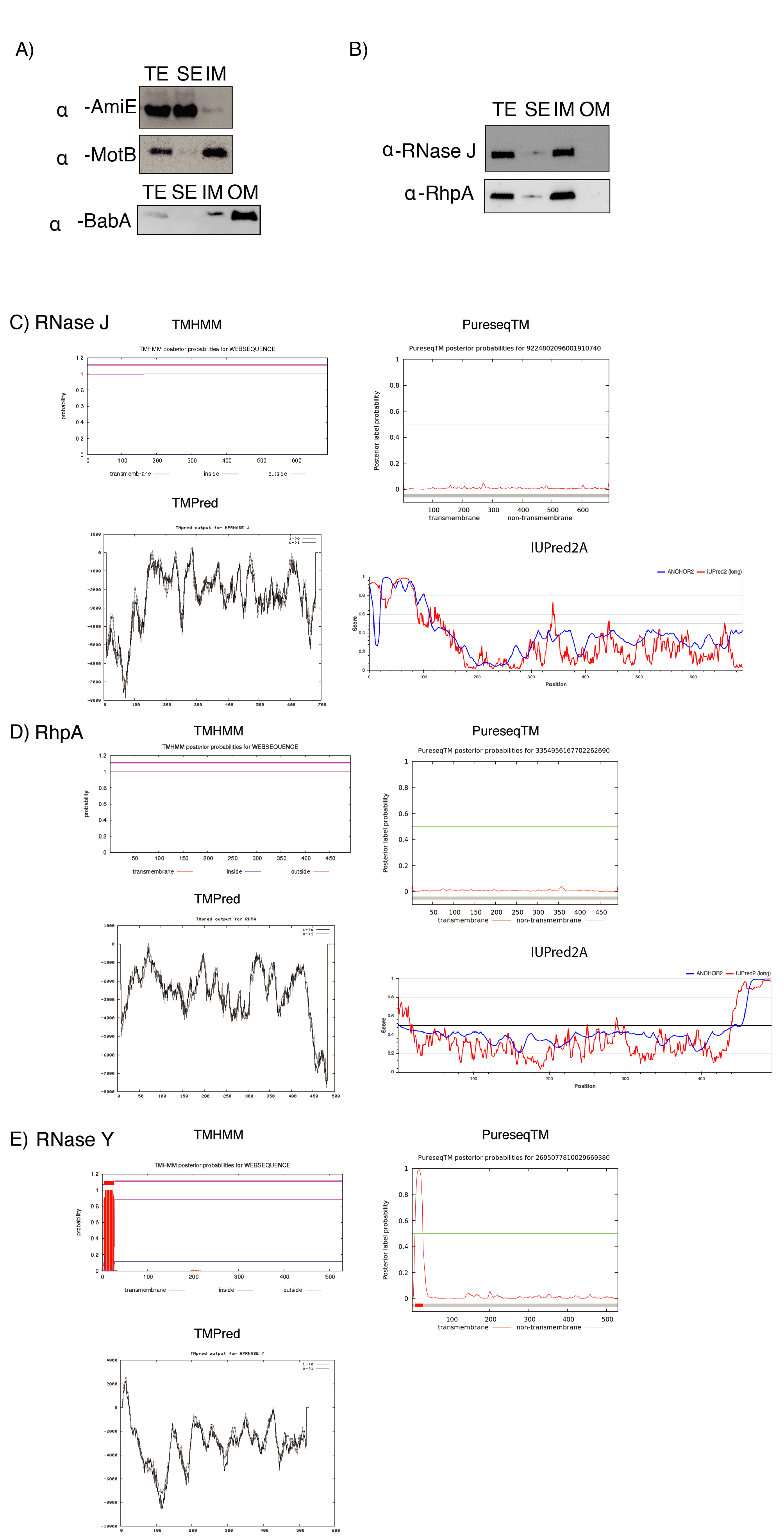

A) Western blots with antibodies against AmiE (soluble extract fraction protein), MotB (inner membrane fraction protein) and BabA (outer membrane fraction protein) on samples submitted to cellular fractionation.

B) Western blots with antibodies against RNase J and RhpA of *H. pylori* strain 26695 on samples submitted to cellular fractionation.

TE: total extract fraction; SE: soluble extract fraction; IM: inner membrane fraction; OM: outer membrane fraction.

C) Prediction of RNase J transmembrane domains by the TMHMM, PureseqTM and TMPred algorithms, and of intrinsically disordered regions (IDRs) by the IUPred2A algorithm.

D) Prediction of RhpA transmembrane and intrinsically disordered regions as in section C.

E) Prediction of RNase Y transmembrane regions as in section C.

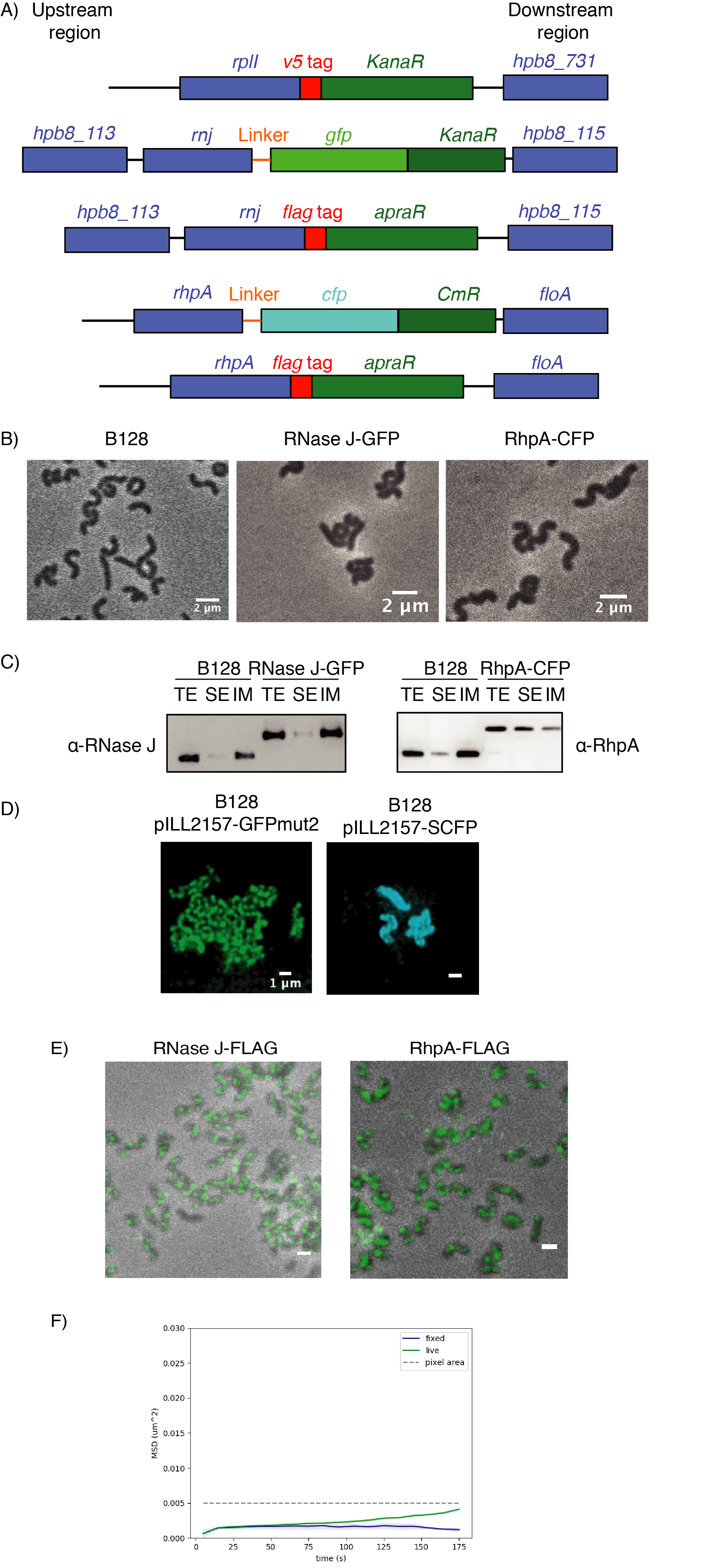

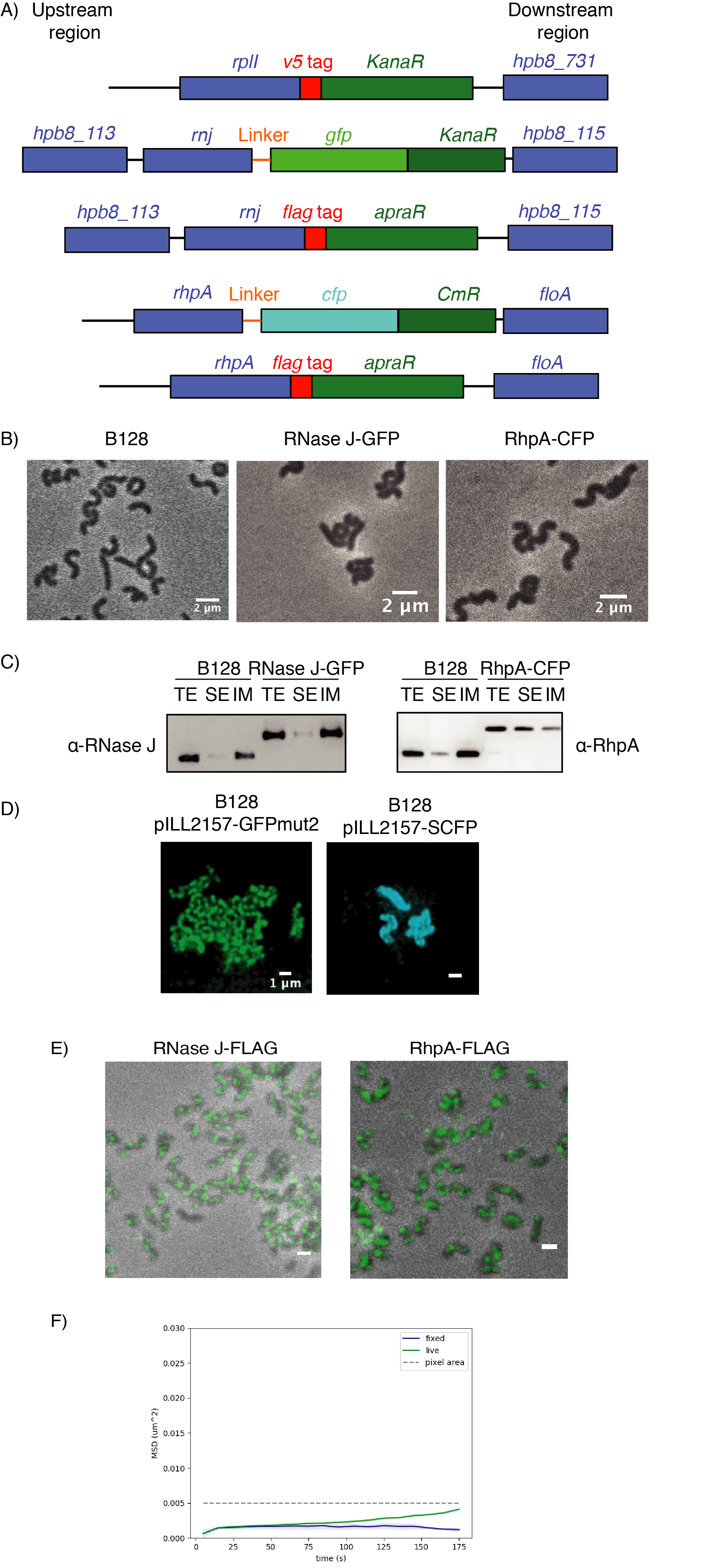

**Figure S2**

A) Schematic representation of the genetic organization of the chromosomal regions of strain B128 expressing L9-V5, RNase J-GFP, RhpA-CFP, RNase J-FLAG and RhpA-FLAG in the corresponding strains. KanaR, kanamycin resistance cassette; ApraR, apramycin resistance cassette; CmR, chloramphenicol resistance cassette.

B) Representative phase contrast microscopy images of B128 wild type and strains expressing RNase J-GFP or RhpA-CFP. No difference in morphology is observed between these strains.

C) Western blots with antibodies against RNase J and RhpA on samples submitted to cellular fractionation of strains expressing or not RNase J-GFP or RhpA-CFP. TE: total extract fraction; SE: soluble extract fraction; IM: inner membrane fraction.

D) Representative confocal fluorescence microscopy images of *H. pylori* B128 strains overexpressing GFP (left) or CFP (right) alone from plasmid pILL2157. Fluorescence is distributed all over the cell.

E) Immunofluorescence confocal microscopy of fixed *H. pylori* cells expressing RNase J-FLAG (left) and RhpA-FLAG (right) with primary antibodies against the FLAG-tag and secondary anti-mice AF488 antibody.

F) The RNase J-GFP foci remain static inside the cells. The displacement of foci was analyzed during 175 min either in live bacteria (n=285 foci) or in fixed bacteria (n=696 foci) as a control. The small change measured in the live bacteria is below the pixel area and thus non-significant. MSD: mean square displacement.

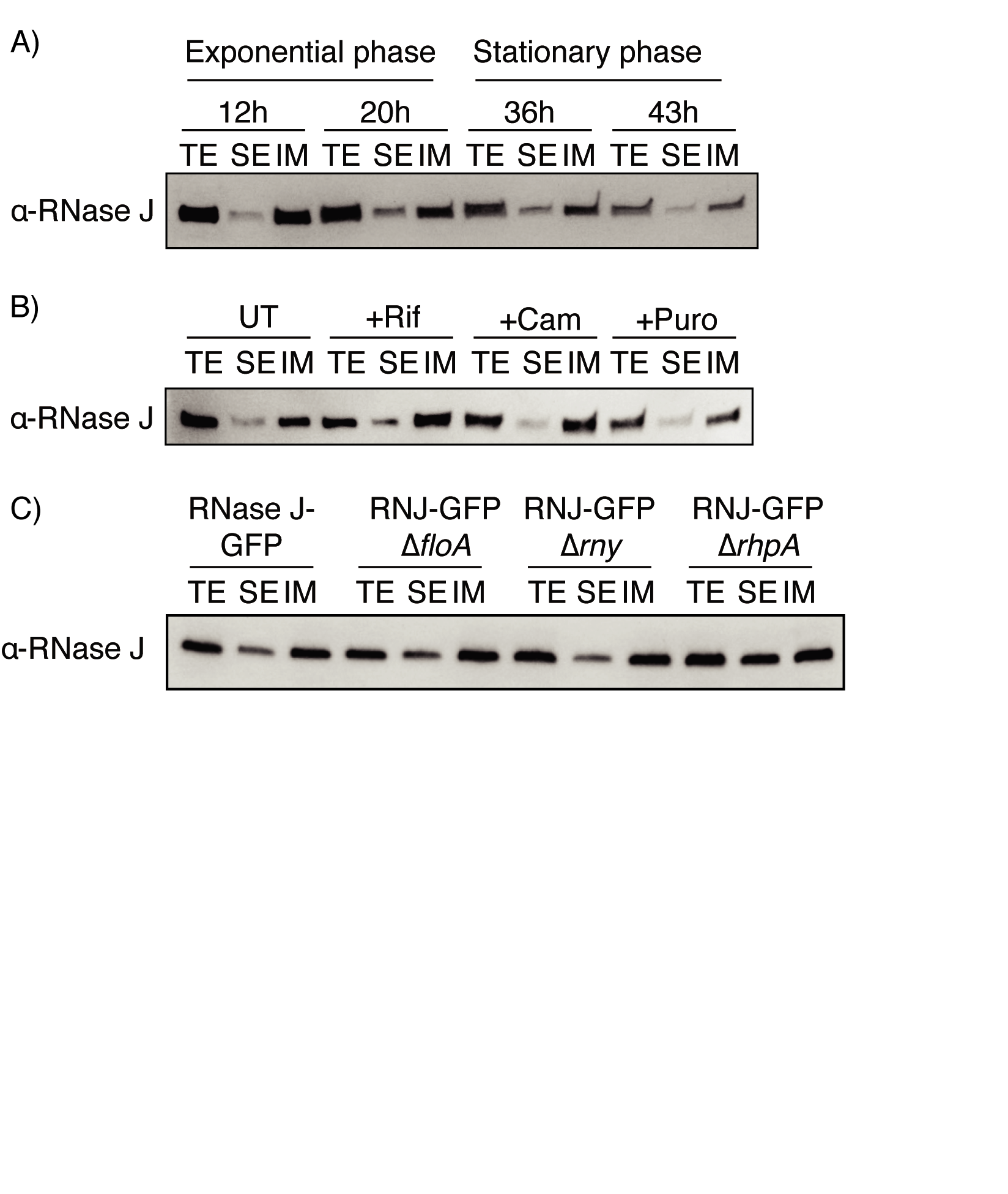

**Figure S3**

The cellular distribution of RNase J-GFP is not affected neither during growth, nor upon antibiotic treatment or in ∆*rny*, ∆*floA* or ∆*rhpA* mutants.

A) Western blot with antibodies against RNase J on samples submitted to fractionation at different timepoints along the growth curve.

B) Western blot with antibodies against RNase J on samples submitted to rifampicin (Rif), chloramphenicol (Cam) or puromycin (Puro) treatment and fractionation.

C) Western blot with antibodies against RNase J on samples from different mutants in exponential phase that were submitted to fractionation. TE: total extract fraction; SE: soluble extract fraction; IM: inner membrane fraction.

**Supplementary video legends**

Supplementary Video 1. Time-lapse of live *H. pylori* expressing RNase J-GFP. The overlay of the phase contrast and GFP channels is shown.

Supplementary Video 2. Time-lapse of fixed *H. pylori* expressing RNase J-GFP. The overlay of the phase contrast and GFP channels is shown.

Supplementary Videos 3-4. 3D visualization of super-resolution imaging data of two representative exponential phase growing bacteria. The yellow-red gradient is the membrane, labeled with WGA-AF555, according to the distance to the coverslip and the blue-green gradient is RNase J-GFP, labeled with anti-GFP-Cy5 nanobodies, according to the distance to the coverslip.

Supplementary Videos 5-6. 3D visualization of super-resolution imaging data of two representative stationary phase growing bacteria. The yellow-red gradient is the membrane, labeled with WGA-AF555, according to the distance to the coverslip and the blue-green gradient is RNase J-GFP, labeled with anti-GFP-Cy5 nanobodies, according to the distance to the coverslip.
