## Supplementary table for "The RNase J-based RNA degradosome is compartmentalized in the gastric pathogen *Helicobacter pylori*"

**Table S1:** Strains, plasmids and oligonucleotides used in this study.

| **Strains** | **Relevant characteristics** | **Reference** |
| --- | --- | --- |
| *Escherichia coli* |  |  |
| DH5α | *F^-^* Φ80*lac*ZΔM15 Δ(*lac*ZYA-*arg*F) U169 *rec*A1 *end*A1 *hsd*R17(r_k_^-^, m_k_^+^) *pho*A *sup*E44 *thi*-1 *gyr*A96 *rel*A1 λ^-^ | (1) |
| OneShot® Top10 | *F-* *mcrA* Δ( *mrr-hsd* RMS-*mcr*BC) Φ80*lac*ZΔM15 Δ*lac*X74 *rec*A1 *ara*D139 Δ(*ara leu*)7697 *gal*U *gal*K *rps*L (StrR) *end*A1 *nup*G | Commercial competent cells (ThermoFisher Scientific) |
| *Helicobacter pylori* |  |  |
| B128 | Sequenced parental strain | (2) |
| B128 ∆*rhpA::apra* | Apra^R^ | (3) |
| B128 *∆rny::apra* | Apra^R^ | This work |
| B128 *∆floA::cm* | Cm^R^ | This work |
| B128 L9-V5 | Km^R^, fusion protein L9-V5 | This work |
| B128 RNase J-GFP | Km^R^, fusion protein RNase J-GFP | This work |
| B128 RNase J-FLAG | Apra^R^, tagged protein RNase J-FLAG | This work |
| B128 RNase J-GFP + ∆*rhpA::apra* | Km^R^, Apra^R^, fusion protein RNase J-GFP and deletion of *rhpA* | This work |
| B128 RNase J-GFP + *∆rny::apra* | Km^R^, Apra^R^, fusion protein RNase J-GFP and deletion of *rny* | This work |
| B128 RNase J-GFP + *∆floA::cm* | Km^R^, Apra^R^, fusion protein RNase J-GFP and deletion of *floA* | This work |
| B128 RhpA-CFP | Cm^R^, fusion protein RhpA-CFP | This work |
| B128 RhpA-FLAG | Apra^R^, tagged protein RhpA-FLAG | This work |
| B128 pILL2157-GFP | Cm^R^, GFP cloned into pILL2157 plasmid | This work |
| B128 pILL2157-CFP | Cm^R^, SCFP cloned into pILL2157 plasmid | This work |

| **Plasmids** | **Relevant characteristics** | **Reference** |
| --- | --- | --- |
| pILL2157 | Expression vector | (4) |
| pILL2157-GFP | pILL2157 containing GFPmut2 | This work |
| pILL2157-CFP | pILL2157 containing SCFP | This work |

References:

1.	Taylor RG, Walker DC, Mclnnes RR. 1993. *E.coli* host strains significantly affect the quality of small scale plasmid DNA preparations used for sequencing. Nucleic Acids Res 21:1677–1678.

2.	McClain MS, Shaffer CL, Israel DA, Peek RM, Cover TL. 2009. Genome sequence analysis of Helicobacter pylori strains associated with gastric ulceration and gastric cancer. BMC Genomics 10:3.

3.	El Mortaji L, Aubert S, Galtier E, Schmitt C, Anger K, Redko Y, Quentin Y, De Reuse H. 2018. The sole DEAD-box RNA helicase of the gastric pathogen Helicobacter pylori is essential for colonization. MBio 9:e02071-17.

4.	Boneca IG, Ecobichon C, Chaput C, Mathieu A, Guadagnini S, Prévost M-C, Colland F, Labigne A, de Reuse H. 2008. Development of inducible systems to engineer conditional mutants of essential genes of Helicobacter pylori. Appl Environ Microbiol 74:2095–102.

| Name |  | Use |
| --- | --- | --- |
| Chromosomal deletions | | |
| oLEM115 | CATTATTCCCTCCAGGTATCAGCCAATCGACTGGCGAG | Rv oligo to amplify the *apraR* cassette. It contains an RBS and an ATG. |
| oLEM120 | TGACTAACTAGGGAGTGCAATGTCGTGCAA | Fw oligo to amplify the *apraR* cassette. It contains a STOP codon upstream of an RBS. |
| oLEM141 | CCATTAGGGCATGAGCTTG | Fw oligo to amplify the region upstream of *rny*. |
| oLEM142 | CATTGCACTCCCTAGTTAGTCAAAGTGGCTTGCCCTCTAGC | Rv oligo to amplify the region upstream of *rny*. It contains a sequence complementary to the *apraR* cassette, a STOP codon and an RBS |
| oLEM143 | GGCAACACGTGGAGCGGATCGGCAATAATCAAGCCTTTTTCC | Fw oligo to amplify the region downstream of *rny*. It contains a sequence complementary to the *apraR* cassette. |
| oLEM144 | GGGTTTGAACGCTTATTGAG | Rv oligo to amplify the region downstream of *rny*. |
| oLEM227 | CAAGGAGGCTGTAATCATC | Fw oligo to amplify upstream region of *rhpA*. |
| oLEM228 | GTATTGCACGACATTGCACTCCGGGAGATTCATACCTCAA | Rv oligo to amplify upstream region of *rhpA*. It contains a homologous region to the *apraR* cassette with its own RBS. |
| oLEM229 | GGCTGATACCTGGAGGGAATAATGCCCATTGATTTGAACG | Fw oligo to amplify downstream region of *rhpA*. It contains a homologous region to the *apraR* cassette with a new RBS. |
| oLEM230 | GCGCACCACAGGGTTGATG | Rv oligo to amplify downstream region of *rhpA*. |
| oATA_001 | TTGCCCCTAAACACCGAGAG | fw primer from 500 bp upstream of *hpb8_1315* (*floA*) |
| oATA_002 | TTTTAGCTTCCTTAGCTCCTGAATTTCCTTTTTAAATATT | rv primer from 20 nt into the chloramphenicol resistance cassette (starting with 2 SD sequences that are included in this primer) |
| oATA_003 | AGGAGCTAAGGAAGCTAAAA | fw primer for the *CmR* cassette (includes the SD sequence at the beginning of that gene) |
| oATA_004 | TTACGCCCCGCCCTGCCACT | rv primer for the chloramphenicol resistance cassette |
| oATA_005 | AGTGGCAGGGCGGGGCGTAATACCTGGAGGGAATAATGAAACGCATGGCGTCTCTTGC | fw primer from the end of *hpb8_1315* (*floA*) (includes 20 last bp from the chloramphenicol resistance cassette, 18 bp that contain a SD sequence to avoid polar effects on *hpb8_1316* and the first 20 bp of the downstream region of that gene) |
| oATA_006 | TAGGGGGCTGTTTTTGAATG | rv primer from the downstream region of *hpb8_1315* (*floA*) |
| Chromosomal fusions | | |
| oEG_003 | AGATGGGTGGTAAGAACAGG | Fw oligo to amplify the end of the *rplI* (L9) gene. |
| oEG_004 | ttaGGTGGAATCCAGGCCCAGCAGCGGGTTCGGGATCGGTTTGCCCTCGGCCACCACATCAATTT | Rv oligo to amplify the end of the *rplI* (L9) gene. It includes a V5 tag. |
| oEG_005 | GGCAAACCGATCCCGAACCCGCTGCTGGGCCTGGATTCCACCtaagaattcgagctcggtacccg | Fw oligo to amplify the v5 tag coupled to a *kanaR* cassette. |
| oLEM010 | CATTATTCCCTCCAGGTAC | Rv oligo to amplify the *kanaR* cassette. |
| oEG_006 | tttagtacctggagggaataATGTTTGAAGCGACAACGAT | Fw oligo to amplify the region downstream of *rplI*. It contains a sequence complementary to the end of the kanaR cassette. |
| oEG_007 | GCAAACGCCTGTAACGATTC | Rv oligo to amplify the region downstream of *rplI*. |
| oLEM196 | CATGGGGAATACAACCATG | Fw oligo to amplify the end of *rnj*. |
| oLEM197 | CATGGATCCATGGAAAAAAGAATGGGCATGACAAATG | Rv oligo to amplify the end of *rnj* without STOP codon. It contains a 12 nt spacer and an ATG for GFP C-terminal fusion. |
| oLEM198 | TTCCATGGATCCATGAGTAAAGGAGAAGAAC | Fw oligo to amplify a 12 nt spacer followed by *gfp*. |
| oLEM199 | GCATGGATGAACTATACAAATGAGGTACCCGGGTGACTAAC | Fw oligo to amplify the *kanaR* cassette. It contains a sequence complementary to the end of *gfp*. |
| oLEM200 | TCATTTGTATAGTTCATCCATGC | Rv oligo to amplify *gfp*. |
| oLEM201 | gtacctggagggaataaTGCCCATTCTTTTtgaTTG | Fw oligo to amplify the end of the *kanaR* cassette and the 16 final nt of *rnj*. |
| oLEM202 | GTGGTAGAAGTCGTTGGAGC | Rv oligo to amplify the region downstream of *rnj*. |
| oATA100 | TTCTTACCCGTGCATGGGG | fw oligo to amplify the last 500 bp of rnj |
| oATA255 | ATTGCACGACATTGCACTCCTTACTTGTCGTCATCGTCTTTGTAGTCAAAAGAATGGGCATGACAAATGATAGCC | rv oligo to amplify the last 500 bp of rnj without STOP codon, including a FLAG tag + STOP codon and a region complementary to the beginning of the apra cassette |
| oATA256 | GGCTGATACCTGGAGGGAATATTGTAACGCTACTGCTATACAAGTCTTAAGAGATGAGGC | fw oligo to amplify the region downstream of rnj with a region complementary to the end of the apra cassette |
| oATA104 | TCGCCTAACGCTAGAGTTAGGG | rv oligo to amplify 500 bp downstream of rnj |
| oATA095 | GGGCTAGACATTAGCGGTGTAAGCC | fw oligo to amplify the last 500 bp of rhpA |
| oATA254 | ATTGCACGACATTGCACTCCTTACTTGTCGTCATCGTCTTTGTAGTCACGGCGTTTGGGTTTTTTAGAATGG | rv oligo to amplify the last 500 bp of *rhpA* without STOP codon, including a FLAG tag + STOP codon and a region complementary to the beginning of the apra cassette |
| oATA097 | CCATTCTAAAAAACCCAAACGCCGTTTCCATGGATCCATGGTGAGCAAGGGCGAGGAG | fw oligo to amplify SCFP from the ATG with the 12 nt spacer at the beginning, for C-terminal fusion + overlapping region with the end of RhpA without STOP codon |
| oATA098 | TTTTAGCTTCCTTAGCTCCTTTACTTGTACAGCTCGTCCATGCCG | rv oligo to amplify SCFP with STOP codon + CamR cassette, including its SD sequence, for C-terminal fusions |
| oATA099 | AGTGGCAGGGCGGGGCGTAAAATATTTAAAAAGGAAATTCATGCCC | fw oligo to amplify the downstream region of RhpA from the STOP codon + overlapping region with the end of CamR with STOP codon |
| Cloning in pILL2157 |  |  |
| gfp-pillA | GTTTGGAAGGAAAAG**caTATG**ATGAGTAAAGGAGAAGAAC | *Nde*I restriction site |
| gfp-pillB | TTTGTATAGTTCATCCATGC |  |
| **gfp**-spa | **GCATGGATGAACTATACAAA**TCCATGGAAAAGAGAAGATG |  |
| spa-pill | GAATTCGAGCTC**GGTACC**CTACTTGTCATCGTCATCC | *Kpn*I restriction site |
| gfp-pill-stop | GAATTCGAGCTC**GGTACC**TTATTTGTATAGTTCATCCATGC | *Kpn*I restriction site |
| oATA019 | **CATATG**GTGAGCAAGGGCGAGGAG | fw primer to amplify *scfp*; contains a *Nde*I restriction site |
| oATA021 | **GGATCC**TTACTTGTACAGCTCGTCC | rv primer to amplify *scfp*; contains a *Bam*HI restriction site |
